# Bacterial inhibition of CD8^+^ T-cells mediated cell death promotes neuroinvasion and within-host persistence

**DOI:** 10.1101/2020.11.20.384974

**Authors:** Claire Maudet, Marouane Kheloufi, Sylvain Levallois, Julien Gaillard, Lei Huang, Charlotte Gaultier, Yu-Huan Tsai, Olivier Disson, Marc Lecuit

## Abstract

Central nervous system infections are amongst the most severe^1,2^, yet the mechanisms by which pathogens access the brain remain poorly understood. The model microorganism *Listeria monocytogenes (Lm)* is a major foodborne pathogen that causes neurolisteriosis, one of the deadliest central nervous system infections^3,4^. While immunosuppression is a well-established host risk factor for neurolisteriosis^3,5^, little is known regarding the bacterial factors underlying *Lm* neuroinvasion. We have developed a clinically-relevant experimental model of neurolisteriosis, using hypervirulent neuroinvasive strains^6^ inoculated in a humanized mouse model of infection^7^, and we show that the bacterial protein InlB protects infected monocytes from CD8^+^ T-cells Fas-mediated cell death, in a c-Met/PI3-kinase/FLIP-dependent manner. This blockade of anti-*Lm* specific cellular immune response lengthens infected monocytes lifespan, favoring *Lm* transfer from infected monocytes to the brain. The intracellular niche created by InlB-mediated cell-autonomous immunosuppression also promotes *Lm* fecal shedding, accounting for its selection as a *Lm* core virulence gene. Here, we have uncovered an unanticipated specific mechanism by which a bacterial pathogen confers to the cells it infects an increased lifespan by rendering them resistant to cell-mediated immunity. This promotes *Lm* within-host persistence and dissemination to the central nervous system, and transmission.

---

**One Sentence Summary:** *Listeria* blocks CD8^+^ T-cells killing and promotes neuroinvasion Bacterial infections of the central nervous system (CNS) are often fatal^1,2^, yet little is known regarding the molecular mechanisms underlying microbial neuroinvasion. *Listeria monocytogenes (Lm)* is a foodborne pathogen that causes neurolisteriosis, one of the deadliest CNS infections^3,4^. Consistent with its key role in immunity against *Lm*, T-cell based immunosuppression is a major host risk factor for neurolisteriosis^3,5^. In contrast, the bacterial factors promoting *Lm* neuroinvasion and their mechanisms of action are poorly understood. Earlier studies have pointed towards the involvement of monocytes in transferring *Lm* from the blood to the brain^8,9^. However, these investigations were performed with reference laboratory *Lm* strains, which are actually poorly neuroinvasive^6^, and require very high bacterial inocula to induce some degree of CNS infection in experimental animal models. This is consistent with the observation that these strains belong to clonal complexes very rarely responsible for human neurolisteriosis^3,6^. In contrast, clinically-associated clonal complexes are hypervirulent and more neuroinvasive^6^. In order to investigate the mechanisms underlying *Lm* neuroinvasion, we developed a clinically-relevant experimental model of neurolisteriosis based on the inoculation of hypervirulent neuroinvasive *Lm* strains^6^ in a humanized mouse model^7^.

## Infected inflammatory monocytes mediate *Lm* neuroinvasion

We inoculated orally *Lm* in humanized KIE16P mice^7^. In contrast to the reference laboratory strain EGDe that belongs to clonal complex (CC) 9 ^10,11^, clinical isolates belonging to the hypervirulent clonal complexes CC1, 4 and 6 systematically induce high-level neuroinvasion, as previously reported^6^, starting at 3 days post-inoculation (dpi) (Fig. 1a). At 5 dpi, the bacterial brain load is the same with or without administration of gentamicin (Fig. 1b), an antibiotic that kills extracellular but not intracellular *Lm* (Extended Data Fig. 1a, b), indicating that intracellular bacteria are involved in neuroinvasion. Consistent with the occurrence of neuroinvasion in the presence of gentamicin, *Lm* is detected intracellularly in the blood and spleen, mainly in inflammatory monocytes (CD45^+^ CD11b^+^ Ly6C^+^ CD3^-^ CD19^-^ CD11c^-^ Ly6G^-^) (Fig. 1c, d and Extended Data Fig. 1c-e), suggesting that monocytes are involved in *Lm* neuroinvasion. This was assessed by infecting *Ccr2^-/-^* mice, in which monocytes are retained in the bone marrow and are therefore less abundant in the blood and spleen^12^ (Extended Data Fig. 1f). Indeed, from day 1 to 3 post-inoculation, more bacteria are gradually recovered in the brain of WT mice as compared to *Ccr2^-/-^* mice (Fig. 1e). Moreover, the transfer of infected monocytes from donor-infected mice into gentamicin-treated uninfected recipient mice is sufficient to induce neuroinvasion, and *Lm* can be recovered from the brain of recipient mice as early as day 2 post-transfer (Fig. 1f). In contrast, the transfer of infected monocytes from mice expressing the diphteria toxin (DT) receptor in myeloid cells (*LysM*-CreER^T2^×iDTR) into recipient mice treated with DT leads to liver and spleen infection but no brain infection, even as late as 4 days post-transfer (Extended Data Fig. 1g, h). Together, these results indicate that infected monocytes are necessary and sufficient to induce neuroinvasion.

**Fig. 1.**
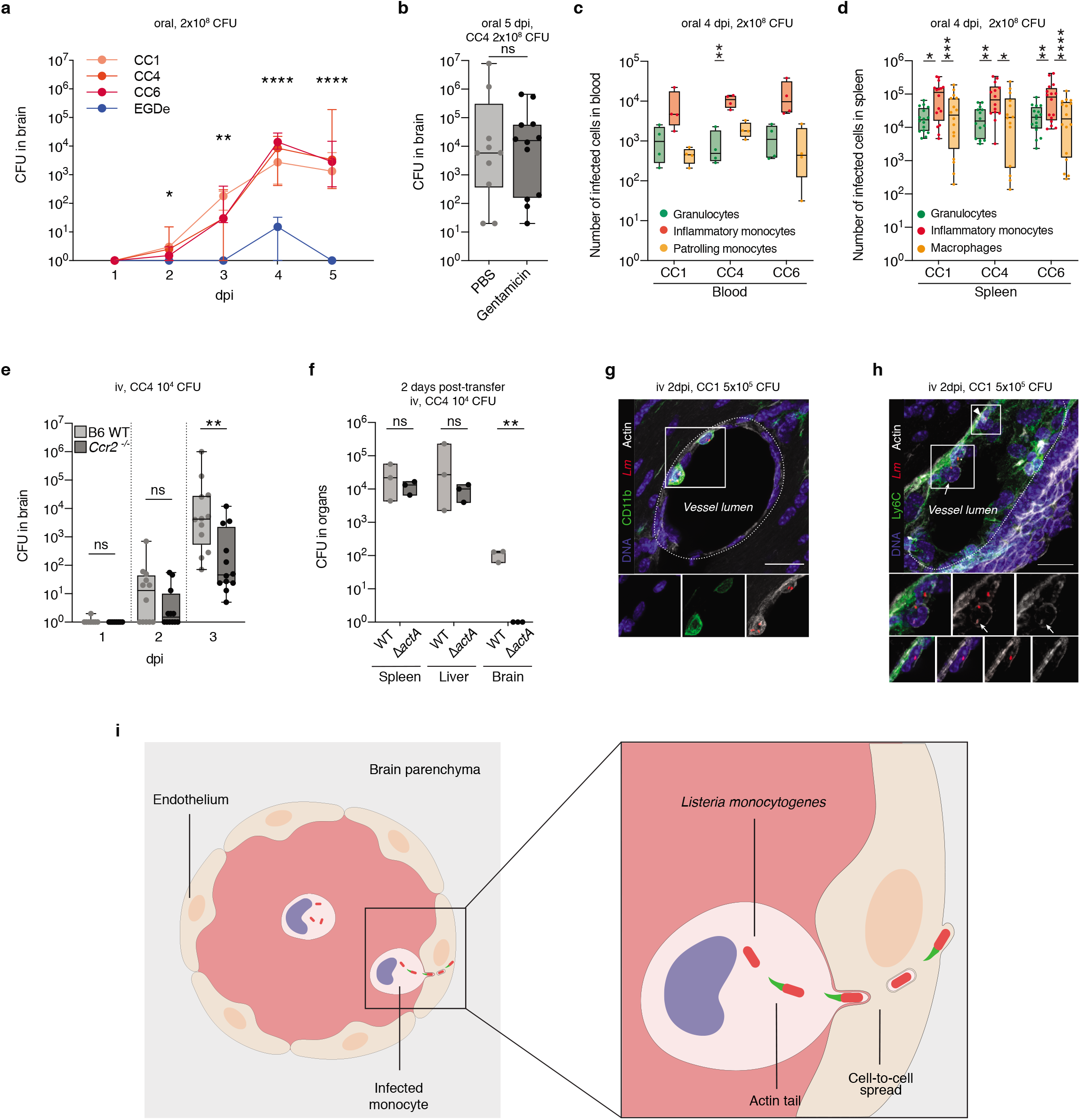
Infected inflammatory monocytes transfer *Lm* to the CNS by cell-to-cell spread. (**a**) Bacterial load in the brain of mice after oral inoculation with CC1/CC4/CC6-*Lm* and EGDe. (**b**) Bacterial load in the brain of gentamicin-treated mice after oral inoculation with *CC4-Lm.* (**c, d**) Three main infected cell subsets in the blood (c) and spleen (d) after oral inoculation with CC1/CC4/CC6-*Lm* in mice at 4 dpi, the peak of bacteremia (Extended Data Fig. 1c). For the blood panel, each dot corresponds to the blood of three mice pooled together. (**e**) Bacterial load in the brain of B6-WT and *Ccr2^-^* mice after intravenous (iv) inoculation with CC4-*Lm*. (**f**) Bacterial load of gentamicin-treated recipient mice, 2 days after injection of infected monocytes harvested from 6 donors mice 3 days after iv inoculation with either CC4-WT or CC4*DactA.* (**g**, **h**) Representative fluorescent microscopy images of infected monocytes adhering to the endothelial cells (g) and of monocytes infected by an actin-polymerizing-*Lm* (arrow) adjacent to an infected endothelial cell (arrowhead, h), 2 days after iv inoculation with CC1-*Lm*. (h) Maximum intensity projection over a 20μm stack and insets are single *z*-planes. Scale bars, 20μm. (**i**) Schematic representation of *Lm* neuroinvasion process. Data were obtained from three (a, b, e and f) and four (c and d) independent experiments and are presented as median ± interquartile (a) and as median ± interquartile (box) and extreme values (lines) (b-f). CFU are compared with the unpaired Mann-Whitney test (a, b, e and f) and number of infected cells with the Friedman test (c and d). ns: *p*>0.05, *:*p* 0.05, **:*p*<0.01, ***:*p*<0.001, ****:*p*<0.0001.

Infected monocytes were observed adhering to the endothelium of blood vessels in brain sections of infected mice (Fig. 1g and Extended Data Fig. 2a-c). In these adhering monocytes, *Lm* polymerizing actin was observed (Extended Data Fig. 2d and movie S1), significantly more than in spleen monocytes (Extended Data Fig. 2e, f), and occasionally adjacent to infected endothelial cells (Fig. 1h and movie S1). Moreover, the transfer of monocytes infected with *LmΔactA* isogenic mutant, unable to polymerize actin and mediate cell-to-cell spread^13,14^, fails to induce neuroinvasion (Fig. 1f). Together, these results demonstrate that *Lm* accesses the brain parenchyma by cell-to-cell spread from adhering bloodborne infected inflammatory monocytes (Fig. 1i). These results are in line with previous reports^8,9^ obtained using poorly-neuroinvasive *Lm* strains, suggesting that neuroinvasive *Lm* strains invade the CNS in a similar manner, albeit to a far greater efficiency (up to 3 orders of magnitude) (Fig 1a).

## InlB increases the number of infected inflammatory monocytes and mediates neuroinvasion

Having identified infected monocytes as critically involved in the onset of neurolisteriosis, we aimed to identify the bacterial factors responsible for *Lm* neuroinvasiveness. Given the well-established role of InlA and InlB in *Lm* crossing of host barriers^7,15^, we investigated their respective role in neuroinvasion. To bypass the well-established role of InlA in the crossing of the intestinal barrier, we inoculated KIE16P mice via the iv route. While InlA plays no role in neuroinvasion (Fig. 2a, Extended Data Fig. 3a-d), InlB plays a major role: the *ΔinlB* mutant is significantly less neuroinvasive than its WT parental strain in co-infection experiments (Fig. 2b, Extended Data Fig. 3a, e). Of note, the *ΔinlAB* mutant is not less neuroinvasive than the *ΔinlB* mutant, ruling out that InlA would act in a conjugated manner with InlB, as it has been reported for placental invasion (Extended Data Fig. 3f, g)^7^. After oral inoculation, a similar difference in neuroinvasiveness is observed between all WT neuroinvasive strains and their corresponding *ΔinlB* mutant (Fig. 2c, Extended Data Fig. 3h). The involvement of InlB in neuroinvasion is also observed upon separate inoculation with either WT-*Lm* or *LmΔinlB* (Fig. 2d, e). InlB contribution to neuroinvasion increases over time (Fig. 2f, g and Extended Data Fig. 3i, j) and *LmΔinlB* never reaches the brain infection level of WT-*Lm* (Fig. 2f, g). These results were confirmed by gene complementation (Extended Data Fig. 3f, g).

**Fig. 2.**
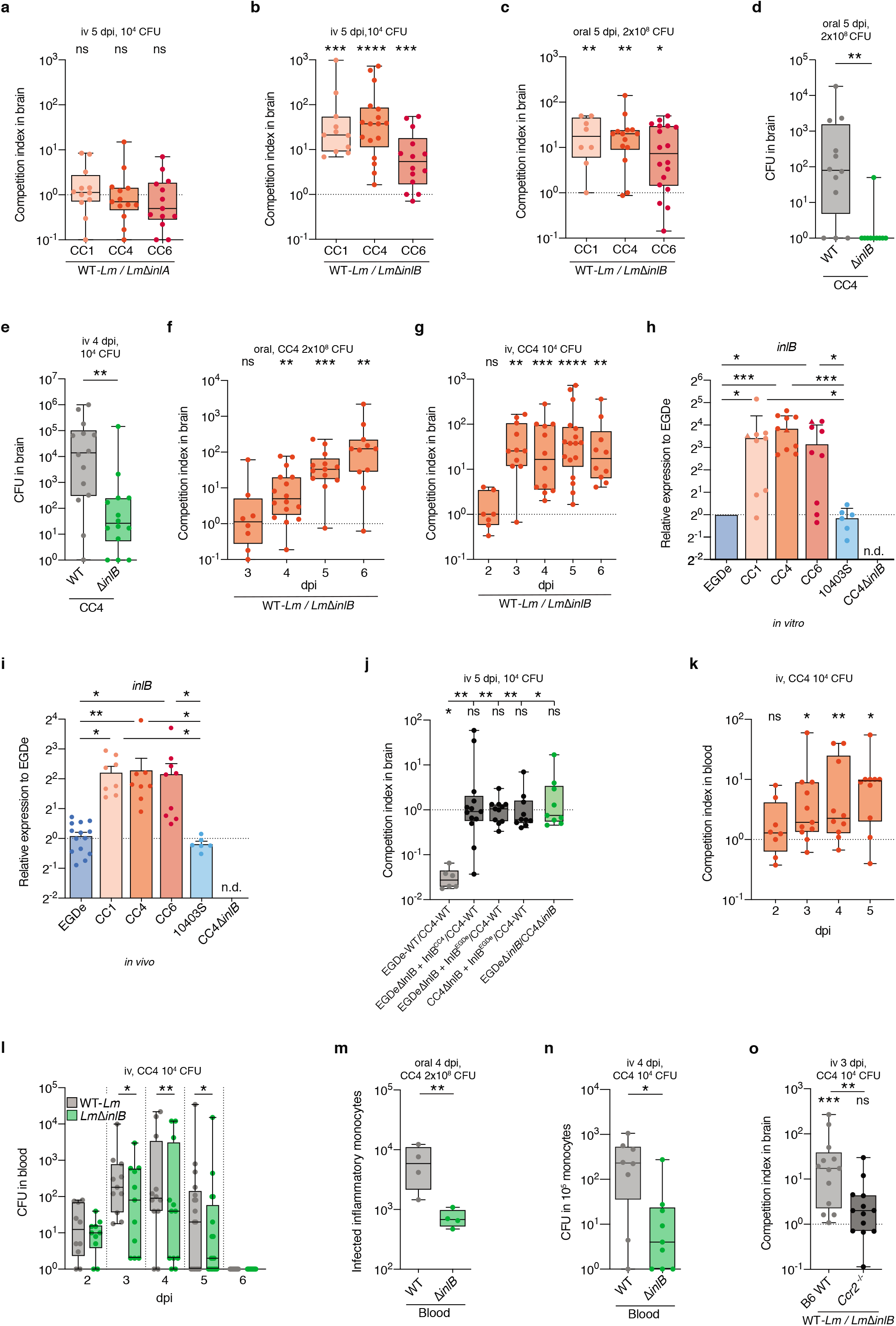
InlB is involved in *Lm* neuroinvasion and inflammatory monocytes infection. (**a-c**) Competition indexes in brain after (a, b) iv and after (c) oral inoculation of WT and isogenic mutant strains. (**d, e**) Bacterial load in brain after (d) oral and (e) iv inoculation with either CC4-WT or *CC4ΔinlB.* (**f, g**) Competition indexes in brain after (f) oral and (g) iv inoculation with 1:1 CC4-WT and CC4Δ*inlB.* (**h, i**) Transcription levels of *inlB* relative to EGDe in (h) mid-log phase in BHI and (i) in infected splenocytes 2 days after iv inoculation. In (h), each dot for CC1/4/6 corresponds to a different clinical isolate and triangles point out the strains used throughout the rest of the study and referred to as CC1, CC4 and CC6. (**j**) Competition index in brain after iv inoculation with a 1:1 mix of the indicated bacterial strains (2×10^4^ for EGDeΔ*inlB* and CC4Δ*inlB*). (**k, l**) Competition index (k) and bacterial load (l) in blood after iv inoculation with 1:1 CC4-WT and *CC4ΔinlB.* (**m, n**) Number of infected monocytes (m) and bacterial enumeration from sorted monocytes (n) in the blood after (m) oral and (n) iv inoculation with CC4-WT or CC4Δ*inlB*. (**o**) Competition index in brain of control or *Ccr2^-/-^* mice after iv inoculation with 1:1 CC4-WT and CC4Δ*inlB*. Data were obtained from three (a-h, j-l and n) and four (i, m and o) independent experiments and are presented as median ± interquartile (box) and extreme values (lines) (a-g and j-o) or as mean ± SD (h and i). CFU in competition assays are compared with the Wilcoxon matched-pairs signed rank test (a-c, f-g, j-l and o) and samples compared with the unpaired Mann-Whitney test (d-e, m-o) or the Kruskal-Wallis test (h-j). ns: *p*>0.05, *: *p*<0.05, **: *p*<0.01, ***: *p*<0.001, ****: *p*<0.0001.

This critical role of *inlB* in *Lm* neuroinvasion was unexpected, as this gene is part of *Lm* core genome and is therefore present in all *Lm* strains, including the poorly neuroinvasive reference strains EGDe and 10403S. Neuroinvasive *Lm* isolates actually strongly upregulate the *inlAB* operon as compared to EGDe and 10403S, both *in vitro* in liquid culture and *in vivo* in infected spleen (Fig 2h, i and Extended Data Fig. 3k-m). To assess the impact of InlB expression level on neuroinvasiveness, we complemented the EGDe*ΔinlB* mutant with the *inlB* gene sequence from either EGDe or CC4 (primary sequences share 93% identity, Extended Data Fig. 4a) so that *inlB* within-host transcription levels are similar to that of endogenous *inlB* in CC4 (Extended Data Fig. 4b). These complemented strains are as neuroinvasive as WT-CC4, whereas CC4*ΔinlB* is as poorly neuroinvasive as EGDe*ΔinlB* (Fig. 2j and Extended Data Fig. c, d). Consistently, CC4*ΔinlB* complemented with the *inlB* EGDe allele expressed to the level of CC4 *in vivo* is as neuroinvasive as WT-CC4 (Fig. 2j and Extended Data Fig. 4 c). Altogether, these results establish that InlB overexpression is the critical factor of *Lm* neuroinvasiveness.

We next evaluated the contribution of InlB to the infection of inflammatory monocytes, having shown their essential role in *Lm* neuroinvasion (Fig. 1). From 3 dpi, the blood bacterial load is higher for WT-*Lm* than *LmΔinlB* (Fig. 2k, l). Moreover, InlB significantly increases the number of *Lm*-infected inflammatory monocytes in the blood and spleen (Fig. 2m, n and Extended Data Fig. 5a, b). In addition, InlB-mediated neuroinvasion is abrogated in *Ccr2^-/-^* mice (Fig. 2o and Extented Data Fig. 5c), indicating that InlB contribution to neuroinvasion implicates infected monocytes. At early time points, when equal numbers of WT and Δ*inlB*-bacteria are retrieved from the blood (1-2 dpi, Extended Data Fig. 5d), equivalent numbers of WT- and *ΔinlB-infected* adhering monocytes are observed in the brain (Extended Data Fig. 2c), showing that InlB does not alter monocyte ability to adhere to brain blood vessels. Moreover, no impact of InlB on bacterial growth is detected upon direct injection of *Lm* into the brain (Extended Data Fig. 5e, f). Altogether, these results indicate that InlB leads to an increased number of circulating infected monocytes, which are themselves required for *Lm* neuroinvasion.

Although InlB has been described as an invasion protein mediating *Lm* internalization into non-phagocytic cells^16,17^, the entry of hypervirulent *Lm* in inflammatory monocytes, which are professional phagocytes, is InlB-independent (Extended Data Fig. 5g-i). This indicates that InlB contribution to neuroinvasion is independent of its capacity to induce internalization.

## InlB-mediated neuroinvasion requires a functional adaptive immune system

T-cell immunosuppression is a well-established risk factor for neurolisteriosis^3,5,18–20^, and hypervirulent *Lm* clones that express the most InlB (Fig. 2h, i) tend to infect patients that are the least immunosuppressed^6^. We therefore hypothesized that InlB may exhibit immunosuppressive properties. Interestingly, treatment with ciclosporin, a prototypic T-cell immunosuppressant, abrogates InlB-dependent neuroinvasion (Fig. 2c, Fig. 3a, b, and Extended Data Fig. 3h), implying that InlB contribution to *Lm* neuroinvasion requires a functional adaptive immunity. Consistently, no difference in neuroinvasion is observed between WT-*Lm* and *LmΔinlB* before 3 to 4 dpi, at a time when adaptive immune responses are not yet expected to be functional^21^ (Fig. 2f, g and Extended Data Fig. 3i, j). Of note, ciclosporin treatment of EGDe-inoculated mice also leads to a slight increased number of infected inflammatory monocytes (Fig. 3c) and increased neuroinvasion (Fig. 3d, e), in line with an immunomodulatory effect of InlB overexpression in hypervirulent isolates. Strikingly, InlB contribution to neuroinvasion is fully abrogated in *Rag2^-/-^* mice (Fig. 3f and Extended Data Fig. 6a-c), that lack functional lymphoid cells, demonstrating their requirement for InlB to exert its immune-mediated effect. The effect of InlB is also abrogated in CD3ε^-/-^, but not in muMt^-/-^ mice (Fig. 3g and Extended Data Fig. 6d-f), and thus depends on T-but not B-lymphocytes.

**Fig. 3.**
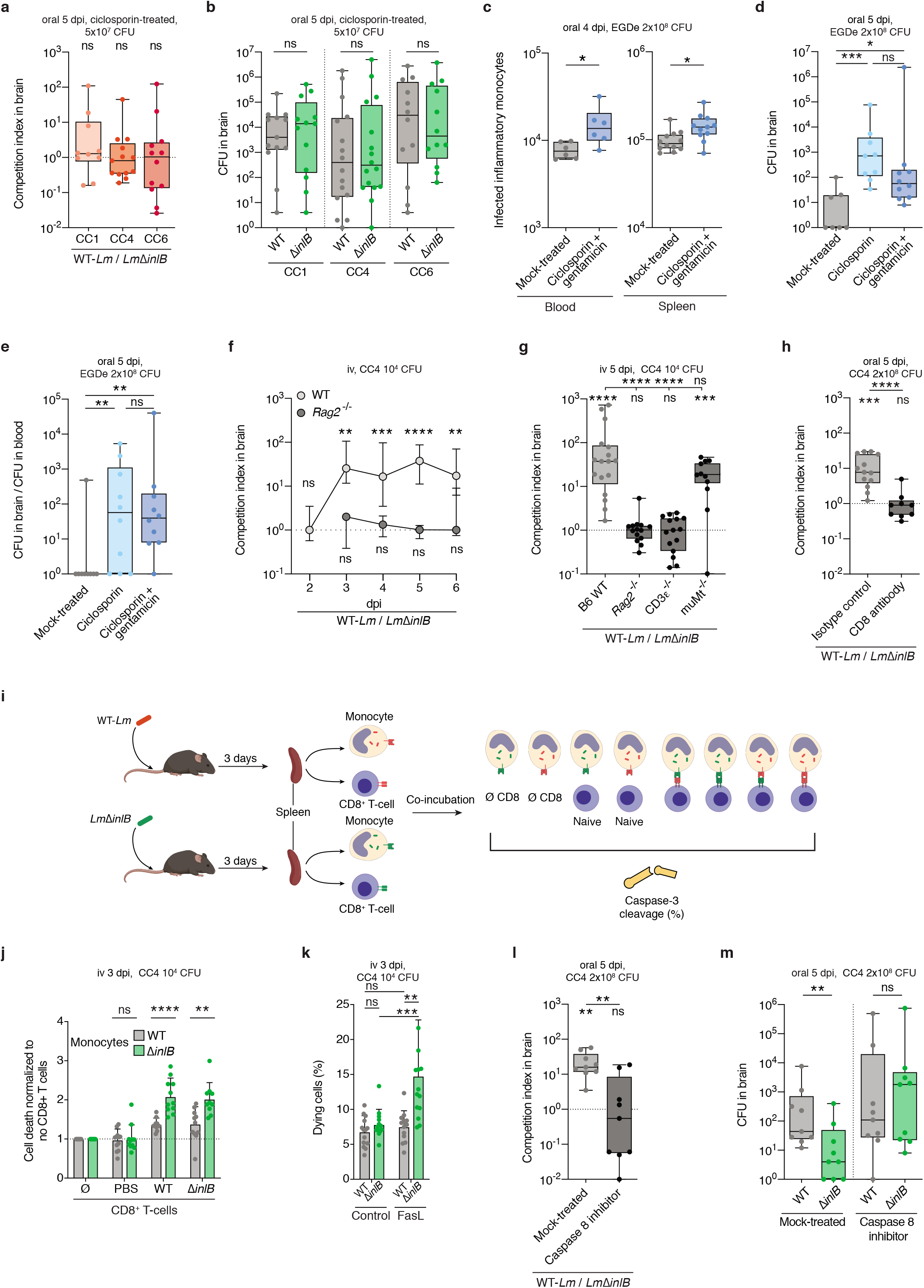
InlB blocks CD8^+^ T-cells-mediated monocyte cell death. (**a, b**) Competition index (a) and bacterial load (b) in brain after inoculation with 1:1 *WT-Lm* strain and *ΔinlB* isogenic mutant in ciclosporin-treated mice, related to Fig. 2c. (**c**) Number of infected monocytes of ciclosporin and gentamicin-treated mice after inoculation with EGDe. (**d, e**) Bacterial load in brain (d) and ratio of brain/blood bacterial load (e) in ciclosporin ± gentamicin-treated mice after inoculation with EGDe. (**f**) Competition index in brain after inoculation with 1:1 CC4-WT strain and *CC4ΔinlB* in control and *Rag2^-/-^* mice. (**g**) Competition index in brain after inoculation with 1:1 CC4-WT strain and *CC4ΔinlB* isogenic mutant in control and in mice lacking functional T (CD3ε^-/-^), B lymphocytes (muMt^-/-^) or both *(Rag2^-/-^).* (**h**) Competition index in brain after inoculation with 1:1 CC4-WT and *CC4ΔinlB* after T-CD8^+^ depletion. (**i**) Schematic pipeline of the cytotoxic lymphocyte (CTL) assay. (**j, k**) Level of caspase-3 cleavage of infected spleen monocytes, harvested after inoculation with CC4-WT or CC4*ΔinlB*, and incubated with CD8^+^ T-cells from similarly infected (WT and *ΔinlB)* or control (PBS) mice at an effector to target ratio of 5 (j) or treated *ex vivo* with FasL (k). (**l, m**) Competition index (l) and bacterial load (m) in the brain after inoculation with 1:1 CC4-WT and CC4Δ*inlB* and after treatment with caspase-8 inhibitor. Data were obtained from three (a-h, l-m) and four (j-k) independent experiments and are presented as median ± interquartile (box) and extreme values (lines) (a-h, l-m) and as mean ± SD (j-k). CFU in competition assays are compared with the Wilcoxon matched-pairs signed rank test (a-b, f-h, l-m), samples are compared with the Mann-Whitney test (c-e, h, l-m), an unpaired student *t*-test (j-k) and the Kruskal-Wallis test (g). ns: *p*>0.05, *: *p*<0.05, **: *p*<0.01, ***:*p*<0.001, ****: *p*<0.0001.

Specifically, CD8^+^ T-cells depletion fully abrogated the effect of InlB (Fig. 3h and Extended Data Fig. 6g, h). Interestingly, specific anti-*Lm* CD8^+^ T-cells are induced to the same extent by WT-*Lm* and *LmΔinlB* isogenic mutant (Extended Data Fig. 7a-d), and accordingly, mice inoculated with WT-*Lm* or *LmΔinlB* display the same level of protective immunity after a second challenge (Extended Data Fig. 7e). Since InlB-mediated neuroinvasion relies on infected monocytes and is detectable in co-infection experiments, we reasoned that InlB may protect specifically infected monocytes from anti-*Lm* T-CD8^+^-mediated specific killing. We therefore performed cytotoxic T-lymphocyte (CTL) assays: infected or uninfected inflammatory monocytes and activated CD8^+^ T-cells were retrieved from mice, either uninfected or infected with WT-*Lm* or *LmΔinlB*, co-incubated and assessed for cell death (Fig. 3i). Strikingly, WT-*Lm*-infected monocytes are protected from CD8^+^ T-cells-mediated cell death, whereas no difference in cell death is observed in uninfected monocytes (Fig. 3j and Extended Data Fig. 8a, b). InlB can be either associated to the bacterial surface or released in the surrounding medium^17^. Consistent with a cell-autonomous effect of InlB, the surface-associated InlB is required for InlB-mediated neuroinvasion (Extended Data Fig. 8c-f). The fact that infection of monocytes is clonal (Extended Data Fig. 8g) and that InlB contribution to neuroinvasion is detectable in co-infection experiments is also fully consistent with the finding that InlB acts in a cell-autonomous manner.

## InlB blocks Fas-mediated cell death, protecting infected cells from killing by CD8^+^ T cells

The cytotoxicity mediated by CD8^+^ T-cells from either *WT-Lm* and *LmΔinlB* infected mice is similar (Fig. 3j), confirming that InlB has no impact on immunization (Extended Data Fig. 7). Since CD8^+^ T-cells cytotoxicity relies on the perforin-granzyme and Fas-Fas ligand pathways^22^, we performed CTL assays in Perforin- and Fas-deficient mice *(Prf1 KO* and *Fas^lpr-cg^*, respectively). InlB-mediated protection against monocyte killing by CD8^+^ T-cells is fully preserved in *Prf1 KO* mice (Extended Data Fig. 8h). In sharp contrast, the inhibitory effect of InlB on infected monocytes cell death is fully abrogated in *Fas^lpr-cg^* mice (Extended Data Fig. 8i). This indicates that InlB blocks Fas-mediated killing whereas the perforin pathway is not involved. Consistently, monocytes infected with WT-*Lm*, but not *LmΔinlB,* are resistant to FasL-induced apoptosis, while surface expression of Fas is not affected by InlB (Fig. 3k and Extended Data Fig. 8j, k). Accordingly, in mice treated with a pharmacological inhibitor of caspase-8, the downstream effector of Fas^23^, *LmΔinlB* becomes as neuroinvasive as WT-*Lm*(Fig. 3l, m). In infected mice, the half-life of monocytes infected by WT-*Lm* is twice longer than that of monocytes infected by *LmΔinlB* (Extended Data Fig. 8l), highlighting that InlB promotes the survival of infected cells. Together, these results demonstrate that InlB blocks CD8^+^ T-cells Fas-mediated killing of infected monocytes.

## InlB inhibition of Fas-mediated CD8^+^ T cell cytotoxicity depends on Met, PI3Kα and FLIP

The receptor of InlB is c-Met^24^, a member of the receptor tyrosine kinases family, which is ubiquitously expressed, including in monocytes^25^. In *Lm*-infected monocytes, bacteria are surrounded by LAMP-1, and InlB induces the recruitment of its receptor c-Met, which can be detected around bacteria (Extended Data Fig. 9a). InlB association to bacterial surface mediates the recruitment of c-Met around bacteria, in contrast to InlB released form, that as expected does not recruits c-Met, and does not mediate neuroinvasion (Extended Data Fig. 8d and 9e). Consistent with a critical role of c-Met, its competitive inhibition by capmatinib fully abrogates InlB-mediated neuroinvasion (Fig. 4a and Extended Data Fig. 9f). In mice where c-Met is conditionally deleted in myeloid cells (*LysM*-CreER^T2^×*Met^flox/+^* or *×Met^flox/flox^)*, InlB-mediated neuroinvasion is also abrogated (Fig. 4b, c). c-Met signals through PI3-kinase (PI3K), leading to the phosphorylation of Akt^26,27^. Consistent with InlB-mediated c-Met engagement in infected monocytes, Akt is phosphorylated in an InlB-dependent manner in these cells (Extended Data Fig. 10a, b). Moreover, inhibition of PI3K activity by the pan-inhibitor wortmannin fully blocks InlB-mediated neuroinvasion (Fig. 4d and Extended Data Fig. 9c). Specifically, inhibition of PI3Kα, but not of the leucocyte-specific PI3Kδ, fully abrogates InlB-mediated neuroinvasion (Fig. 4e and Extended Data Fig. 10d, e). FLIP, a PI3K-regulated competitive inhibitor of procaspase-8^28^,^29^, is upregulated in infected monocytes, in an InlB-dependent manner, resulting in a decreased activity of caspase-8 (Fig. 4f, g and Extended Data Fig. 10f). Inhibition of either c-Met or PI3Kα blocks both InlB-mediated FLIP upregulation and the decrease of caspase-8 activity (Fig. 4f, g). In mice conditionally deleted for FLIP, InlB-mediated monocytes resistance to cell death is lost (Fig. 4h and Extended Data Fig. 10g). Moreover, in mice where FLIP is conditionally deleted in myeloid cells (*LysM-*CreER^T2^×*FLIP^flox/+^* or *×FLIP^flox/flox^)*, InlB involvement in neuroinvasion is fully abrogated (Fig. 4i, j). Altogether, these results demonstrate that InlB-mediated blockade of the Fas cell death pathway in infected monocytes results from the PI3Kα-dependent cell autonomous upregulation of the caspase-8 inhibitor FLIP (Extended Data Fig. 10h).

**Fig. 4.**
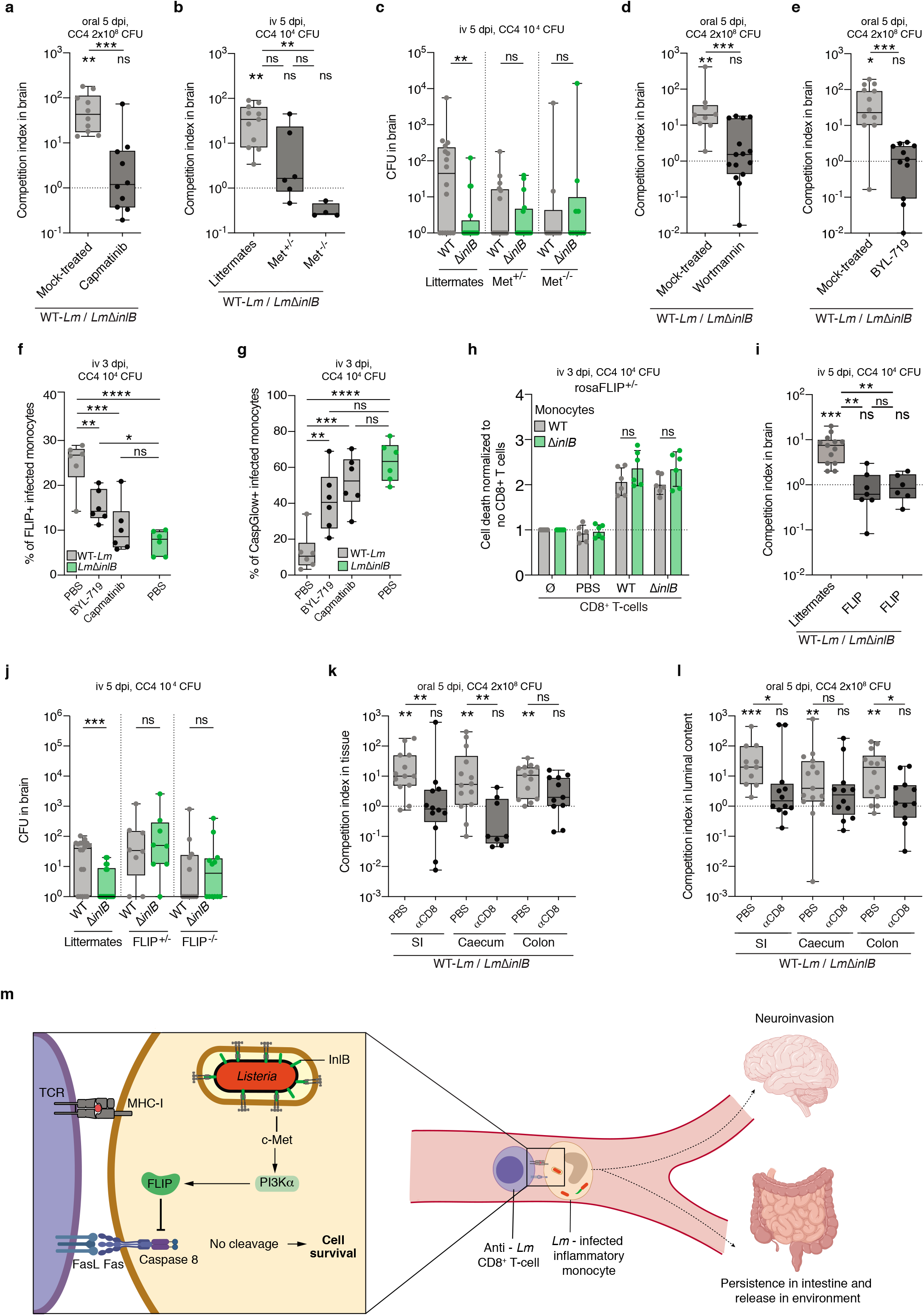
Inhibition of CD8^+^ T cells-mediated cell death by InlB is dependent on the c-Met-PI3K α-FLIP pathway and is involved in *Lm* intestinal colonization and fecal carriage. (**a**) Competition index in the brain after inoculation with 1:1 of CC4-WT and *CC4ΔinlB,* in mice treated with a c-Met inhibitor (capmatinib). (**b, c**) Competition index (b) and bacterial load (c) in the brain of *LysM*-CreER^T2^ x *Met*^flox/flox^ (or *Met*^+/flox^) mice (referred to as Met^-/-^ and Met^+/-^ mice), and their littermates, after inoculation with 1:1 of CC4-WT and CC4Δ*inlB* and tamoxifen treatment. (**d, e**) Competition indexes in the brain after inoculation with 1:1 of CC4-WT and *CC4ΔinlB,* in mice treated with a pan-PI3K inhibitor (wortmannin, d) or a specific PI3Kαinhibitor (BYL-719, e). (**f, g**) Proportion of infected monocytes expressing FLIP (f) or active caspase-8 (g) after inoculation with CC4-WT and *CC4ΔinlB* in mice treated with either BYL-719 or capmatinib. (**h**) Level of caspase-3 cleavage of infected spleen monocytes, harvested after inoculation with CC4-WT or *CC4ΔinlB* of tamoxifen-treated *Rosa26-CreER^T2^* × *Cflar*^+/flox^ (rosaFLIP^+/-^) mice and incubated with CD8^+^ T cells from similarly infected mice at an effector to target ratio of 5. (**i, j**) Competition index (i) and bacterial load (j) in *LysM*-CreER^T2^ × FLIP^flox/flox^ (or FLIP^+/flox^) mice (referred to as FLIP^-/-^ and FLIP^+/-^ mice), and their littermates, after inoculation with 1:1 of CC4-WT and CC4Δ*inlB* and tamoxifen treatment. (**k, l**) Competition index in the intestinal tissues (k) and content (l) after inoculation with 1:1 CC4-WT and *CC4ΔinlB* in mice treated with an anti-CD8^+^ T-cells antibody, SI = small intestine. (**m**) Schematic representation of InlB-mediated blockade of CD8+ T cell-mediated cell death resulting in neuroinvasion, persistence in the intestine and transmission. Data were obtained from three (f-h, k-l) or four (a-e, i-j) independent experiments and are presented as median ± interquartile (box) and extreme values (lines) (a-g, i-l) or mean ± SD (h). CFU in competition assays are compared with the Wilcoxon matched-pairs signed rank test (a-e, i-l) and samples are compared with the Mann-Whitney test (a, d-e, k-1), an unpaired student t-test (h), a Kruskal-Wallis test (b and i) and a one-way ANOVA (f-g). ns: *p*>0.05, *: *p*<0.05, **: *p*<0.01, ***: *p*<0.001, ****: *p*<0.0001.

## InlB inhibition of CD8^+^ T cell mediated killing promotes *Lm* persistence and fecal carriage

InlB is part of *Lm* core genome and is under purifying selection^30,31^, suggesting that InlB confers a selective advantage to *Lm*. As *Lm* is shed back from infected tissues into the intestinal lumen^32^, *Lm* increased virulence may translate into increased fecal shedding, thereby favoring transmission. We therefore tested whether InlB is also involved in *Lm* intestinal carriage and release in the feces. Indeed, WT-*Lm* levels of infection of intestinal tissues and release in the intestinal lumen and feces are significantly higher than that of *LmΔinlB,* and these differences, as for neuroinvasion, are fully dependent on CD8^+^ T-cells (Fig. 4k, l). These observations highlight that InlB-mediated neuroinvasion, although resulting from a very specific interference with adaptive immunity, likely reflects the selective advantage provided by InlB to *Lm* for within-host persistence and inter-host transmission (Fig 4m).

## Discussion

Here we uncovered that *Lm* is able to render its host cell resistant to CD8^+^ T-cells-mediated killing. Selective blockade of the Fas-FasL death pathway in infected monocytes allows these cells to survive longer in the blood, and ultimately transfer *Lm* more abundantly to the brain. Importantly, this novel, specific and unanticipated mechanism that creates an intracellular protected niche for *Lm* is also involved in its persistence in the intestinal tissue and release in the environment. This mechanism is mediated by the surface protein InlB, which was so far described as involved in *Lm* internalization into non-phagocytic cells, whereas we uncover here its key role as an immunomodulatory protein. InlB promotes monocytes survival and neuroinvasion through a refined and unsuspected mechanism: the upregulation, *via* a c-Met/PI3Kα-dependent pathway, of the anti-apoptotic factor FLIP, which competitively inhibits caspase-8 cleavage and blocks Fas-FasL mediated cell death.

This study highlights the critical role played by cellular immunity against intracellular pathogens’ neuroinvasion from a microbial perspective. Indeed, whereas extracellular pathogens rely on the binding to specific host cell receptors^33^ or the breaching of host barriers^34^ to invade the CNS, the facultative intracellular pathogen *Lm* takes advantage of its ability to invade and persist within host cells to access to the brain. *Lm* intracellular persistence mediated by InlB promotes *Lm* neuroinvasion *via* ActA-dependent transfer of *Lm* from infected monocytes to brain endothelial cells. These results show that the capacity of microbes to survive into cells is a key pathogenic determinant favoring within-host dissemination and ultimately neuroinvasion. Other neuroinvasive intracellular pathogens, such as *Mycobacterium tuberculosis* and *Toxoplasma gondii,* also stimulate PI3K^35,36^ and survive in myeloid cells. As *Lm*, these neuroinvasive microorganisms may also protect infected cells from killing by the immune system, favoring their survival, and increasing their within-host persistence and neuroinvasiveness.

Intracellular pathogens tend to interfere with innate immune responses to establish a successful infection, and some also interfere with the adaptive immune system in a broad and non-selective manner^37,38^, favoring their persistence, like HIV, EBV and measles virus. *Lm* has been instrumental for the discovery of cellular immunity^39^ and is indeed a prototypic inducer of a protective CD8^+^ T-cell response^20,40^. Yet, we have shown that InlB selectively blocks the action of the most efficient and specific anti-*Lm* immune effector, T cells-mediated cytotoxicity. This allows the establishment of a protected niche favoring *Lm* dissemination and persistence within the host. This is reminiscent of the mechanism by which tumor cells, in which signaling downstream of growth factor receptors is frequently constitutively activated^41,42^, also evade immune responses by surviving immune killing. A detailed understanding of how microbes have selected mechanisms to interfere with the immune system may help to rationally design novel anti-infective and anti-tumor therapies. Similarly, the immunomodulatory mechanism of InlB, specific of and restricted to infected cells, may also help develop new immunosuppressive therapies aimed at specifically protecting cells of interest from the immune system, as opposed to classic immunosuppressive drugs that inhibit indiscriminately immune functions, and therefore favor infectious and neoplastic complications.

*Lm* is an opportunistic pathogen that only rarely induces clinically apparent infection upon oral ingestion^43^, and there is no inter-human transmission of listeriosis. Yet, the so-called “virulence factors” of *Lm* are under purifying selection^30,31,44^, implying that they contribute to *Lm* fitness. By interfering with the host anti-*Lm* cellular effectors, we have shown that InlB enhances *Lm* intestinal carriage and fecal shedding, thereby increasing the odds of neuroinvasive *Lm* to be transmitted back to the environment and colonize new hosts. This illustrates that the anthropocentric view on microbial pathogenesis which phenotypic output is centered on disease does not necessarily reflect the actual context where microbial evolution and fitness gain take place.

## Acknowledgments

We thank Philippe Bousso and Alain Fischer for helpful discussions, and the members of the Biology of Infection Unit for their support, in particular Laetitia Travier for technical help on brain microscopy and Lukas Hafner for contributing to data analysis. We thank the Cytometry and Biomarkers Unit of Technology and Service (CB UTechS), Dmity Ershov from the Image Analysis Hub and the Center for Animal Resources and Research (C2RA) at Institut Pasteur. We are grateful to Geneviève de Saint Basile and Fernando Sepuvelda (Institut Imagine, Paris) for the Prf1 KO mice, Frédéric Rieux-Laucat (Institut Imagine, Paris) for the Fas^lpr-cg^ mice, Richard Pope for the FLIP^flox/flox^ mice, Florian Greten for the LysMCreER^T2^ mice, Alain Eychene for the Met^flox/flox^ mice, and Javier Pizarro-Cerda and Pascale Cossart (Institut Pasteur) for the pAD β-lactamase plasmid.

## Funding

Work in ML laboratory is funded by Institut Pasteur, Inserm, ERC, ANR and Labex IBEID (ANR-10-LABX-62-IBEID). CM was a recipient of the Roux-Cantarini fellowship of Institut Pasteur. LH, CG and JG were supported by Université Paris Descartes, YHT by the Pasteur - Paris University (PPU) International PhD Program, under the European Union’s Horizon 2020 research and innovation program, Marie Sklodowska-Curie grant agreement No 665807, and SL by FRM (ECO201906009119) and “Ecole Doctorale FIRE-Programme Bettencourt”.

## Author contributions

CM, MK, and ML designed the experimental strategy. CM, MK, SL, JG, YHT and OD designed and performed mouse experiments. CM, LH and CG designed and performed *in vitro* experiments. CM, MK, SL, JG, OD, YHT and ML analyzed the data. CM, SL and ML wrote the manuscript, MK and OD edited it, and all authors agreed on its final version.

## Competing interests

The authors declare no competing interests.

## Data and material availability

The datasets generated during and/or analyzed during the current study are available from the corresponding author on reasonable request.

## Extended Data

Materials and Methods

Extended Data Tables 1-5

Extended Data Figures 1-10

Extended Data Movie 1

Extended References (45-60)

---

**Extended Data Fig. 1.**
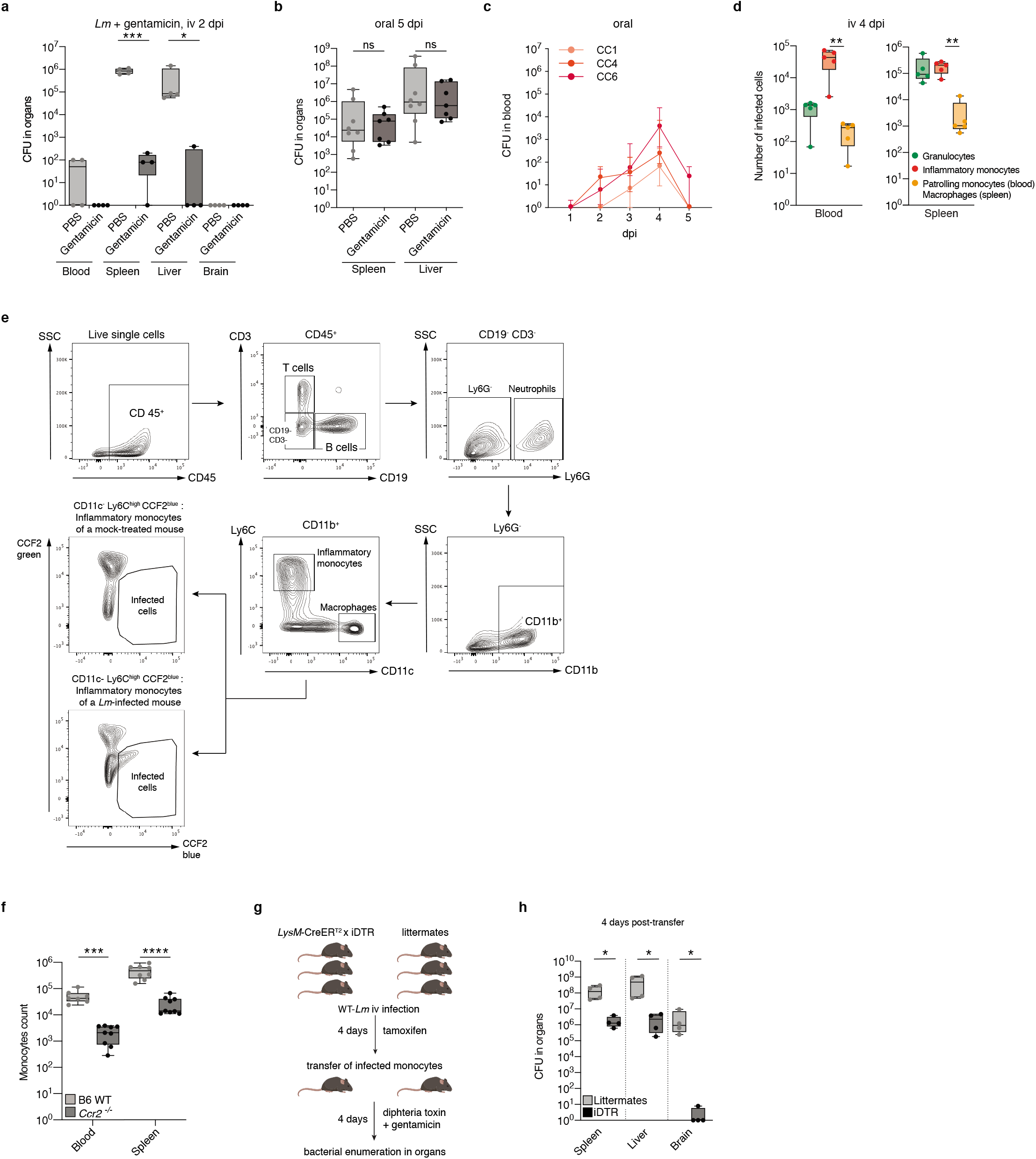
Infected inflammatory monocytes are necessary for *Lm* to invade the CNS. (**a**) Bacterial load in organs of KIE16P mice 2 days after iv inoculation with 10^4^ CFU CC4-*Lm* immediately followed by injection of gentamicin, assessing the bactericidal effect of gentamicin on extracellular circulating *Lm*. (**b**) Bacterial load in the spleen and liver 5 days after oral inoculation with 2×10^8^ CFU CC4-*Lm*, in KIE16P mice treated with gentamicin every day from day 1 post-inoculation, related to Fig. 1A. (**c**) Bacterial load in the blood of KIE16P mice after oral inoculation with 2×10^8^ CFU of CC1/CC4/CC6-*Lm*, related to Fig. 1A. (**d**) Repartition of the 3 main infected cell subsets in the blood and spleen of KIE16P mice 4 days after iv inoculation with 10^4^ CFU of CC4-*Lm*. (**e**) Representative dot plots of the gating strategy used for flow cytometry analysis. Infected cells are identified through the shift of fluorescence, upon excitation with the 405 nm laser, of the CCF2-AM substrate from green (518 nm) to blue (447 nm) in presence of β-lactamase expressing-*Lm*. (**f**) Number of inflammatory monocytes in the blood and spleen of B6-WT or *Ccr2^-/-^* mice. (**g**) Schematic pipeline of the transfer experiment in *LysM-*CreER^T2^× iDTR mice. (**h**) Bacterial load in the spleen, liver and brain of gentamicin- and diphtheria toxin-treated recipient *LysM*-CreER^T2+/-^×*Rosa26*-iDTR^+/-^ and littermates mice, 4 days after injection of infected monocytes harvested from *n*= 3 donor tamoxifen-treated *LysM-*CreER^T2+/-^×*Rosa*26-iDTR^+/-^ or *n*= 3 littermates mice, 4 days after iv inoculation with 10^4^ CC4-WT. Data were obtained from two (a) or three (b-d, f) and four (h) independent experiments and are presented as median ± interquartile (box) and extreme values (lines) (a-b, d, f and h) or as median ± interquartile (c). Samples are compared with an unpaired Mann-Whitney test (a-b, f and h) and number of infected cells with the Friedman test (d). ns:*p*>0.05, *:*p*<0.05, **:*p*<0.01, ***: *p*<0,001, ****: *p*<0.0001.

**Extended Data Fig. 2.**
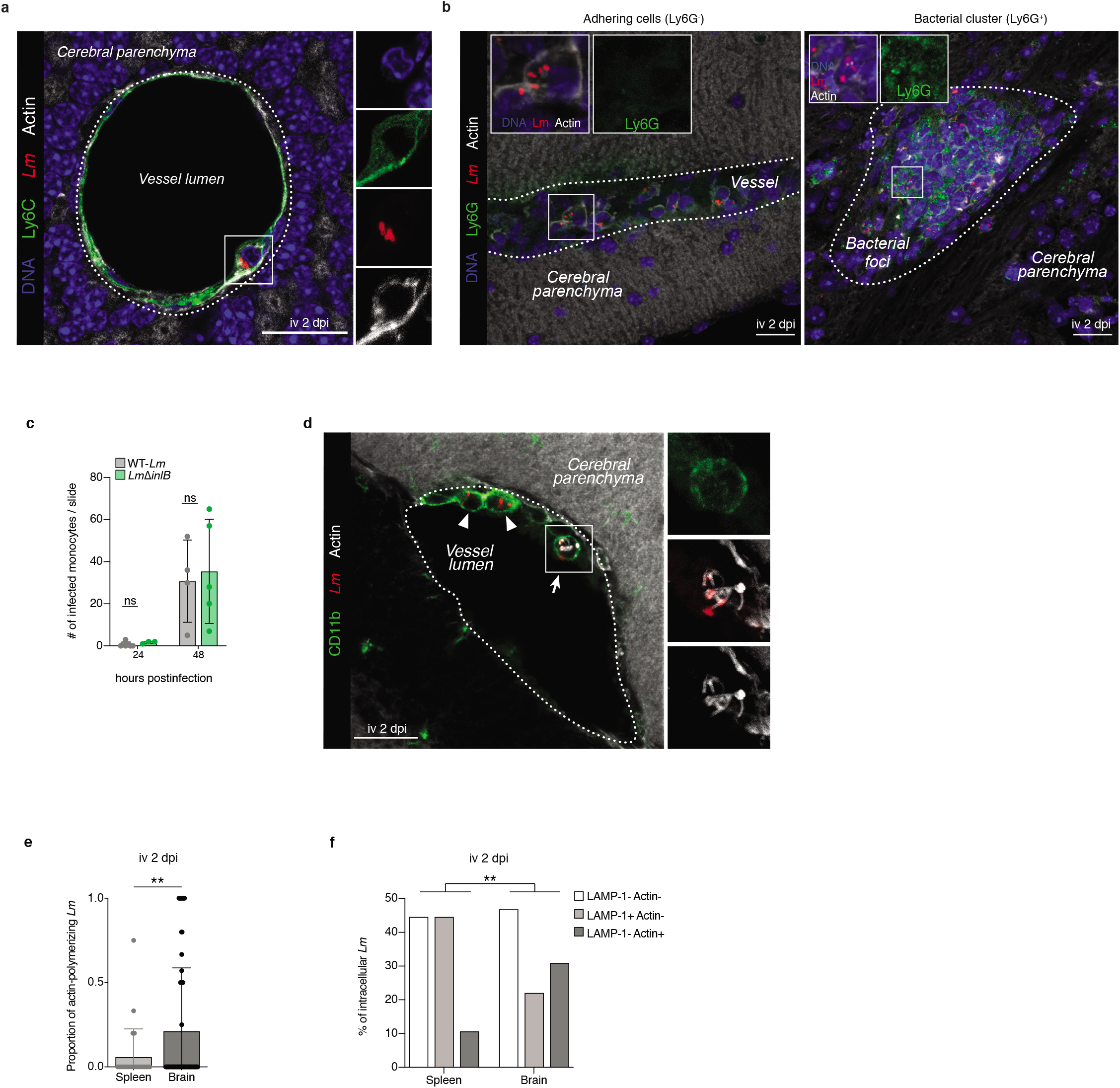
Infected inflammatory monocytes transfer *Lm* to the CNS. (**a-b**) Representative fluorescence microscopy images of infected inflammatory monocytes adhering to endothelial cells 2 days after iv inoculation with 5×10^5^ CFU *CC1-Lm* in KIE16P mice. Adhering infected cells are Ly6C^+^ (a) and Ly6G^-^ (b left panel; to ensure the specificity of the Ly6G^-^ staining we show in the right panel a positive control staining for Ly6G in a parenchymal bacterial cluster). **(c)** Quantification of infected monocytes in brain vessels of KIE16P mice 24 and 48 hours after iv inoculation with 5×10^5^ CFU CC4-*Lm* and *CC4ΔinlB.* Each dot corresponds to the average number of monocytes counted on two slides (representative median sagittal sections, 40 μm thickness) for one mouse. **(d)** Representative fluorescence microscopy images of infected inflammatory monocytes adhering to endothelial cells 2 days after iv inoculation of 5×10^5^ CFU CC1-*Lm* in KIE16P mice, in which intra-monocytic *Lm* are found polymerizing host actin. **(e, f)** Proportion of actin-polymerizing *Lm* (e) and *Lm* co-localizing with either LAMP-1 or Actin (f) in spleen and brain of KIE16P mice 3 days after iv inoculation with 10^4^ CC4-*Lm*. Scale bars: 20 μm, a and d are maximum intensity projections over a *z*-stack. Data were obtained from three independent experiments and are presented as mean ± SD. Samples are compared with an unpaired Mann-Whitney test (c and e) and a one-way ANOVA (f). ns: *p*>0.05, **: *p*<0.01.

**Extended Data Fig. 3.**
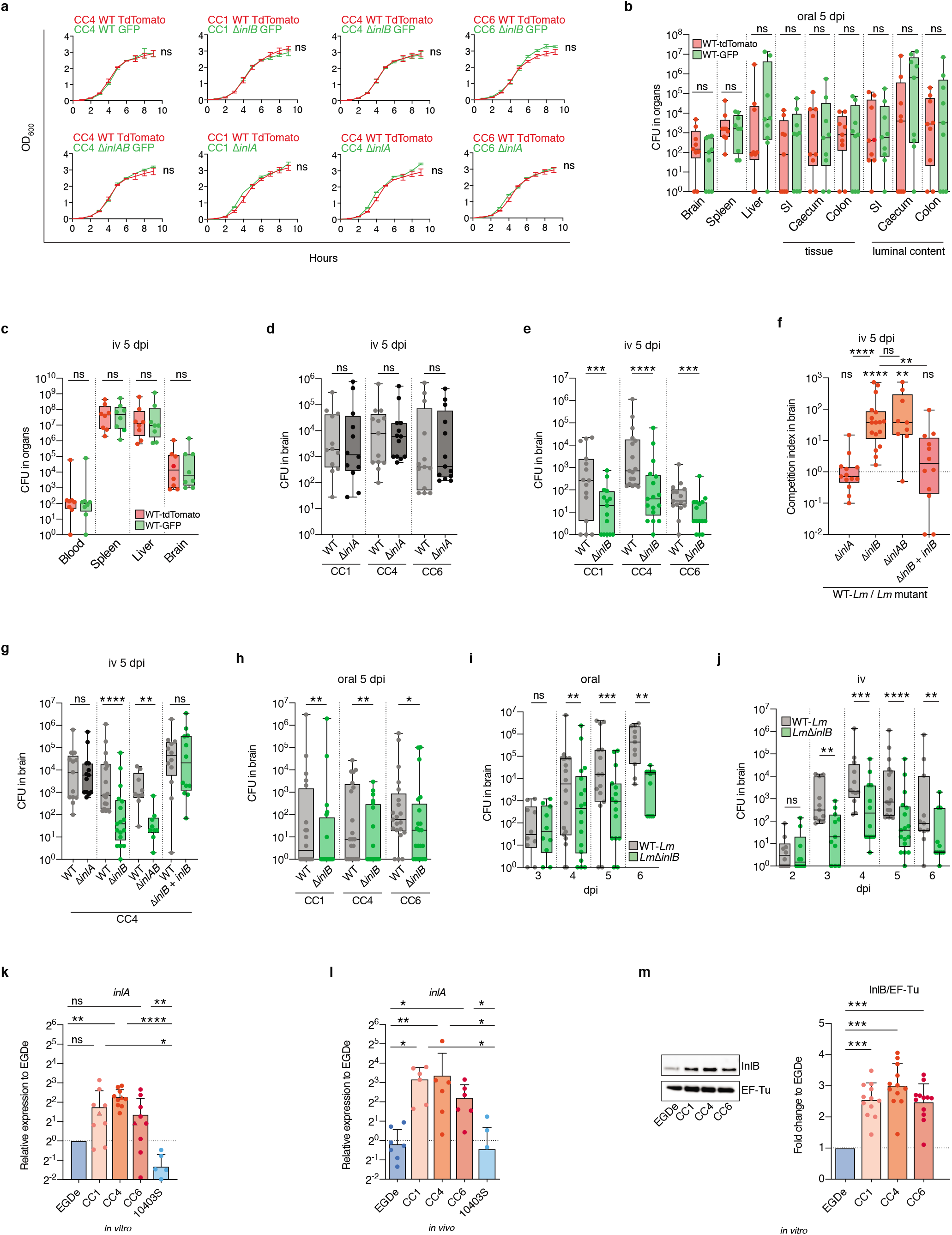
InlB is a major determinant of *Lm* neuroinvasiveness whereas InlA is not. **(a)** Optical density of indicated bacterial strains measured every hour for 9 hours after 1:100 dilution in BHI of an overnight culture. (**b** and **c**) Bacterial load in organs or luminal contents of KIE16P mice 5 days after oral inoculation with 2×10^8^ CFU (b) or after iv inoculation with 10^4^CFU (c) of 1:1 CC4-WT expressing TdTomato or GFP. **(d)** Bacterial load in brain of KIE16P mice 5 days after iv inoculation with 10^4^ CFU of 1:1 mix of WT and *ΔinlA* isogenic strains, related to Fig. 2a. **(e)** Bacterial load in brain of KIE16P mice 5 days after iv inoculation with 10^4^ CFU of 1:1 mix of WT and *ΔinlB* isogenic strains, related to Fig. 2b. **(f, g)** Competition index (f) and bacterial load (g) in brain of KIE16P mice 5 days after iv inoculation with 10^4^ CFU of 1:1 mix of CC4-WT and either *CC4ΔinlA, CC4ΔinlB, CC4ΔinlAB* or *CC4ΔinlB* complemented with *inlB,* related to Fig. 2a-b and panels d,e. **(h)** Bacterial load in brain of KIE16P mice 5 days after oral inoculation with 2×10^8^ CFU of 1:1 mix of WT strain and *ΔinlB* isogenic strains, related to Fig. 2c. **(i, j)** Bacterial load in brain of KIE16P mice at indicated times after oral inoculation with 2×10^8^ CFU (i) and after iv inoculation with 10^4^ CFU (j) of 1:1 CC4-WT and CC4Δ*inlB*, related to Fig. 2f, g. **(k)** Transcription levels of *inlA* relative to EGDe in mid-log phase in BHI. For CC1/4/6, each dot corresponds to a different clinical isolate and triangles represent the strains used throughout the rest of the study and referred to as CC1, CC4 and CC6, related to Fig. 2h. **(l)** Transcription levels of *inlA* relative to EGDe in infected splenocytes 2 days after iv inoculation with 2×10^5^ CFU in KIE16P mice, related to Fig. 2i. **(m)** Representative Western blot (left) and quantification (right) of InlB expression, normalized to that of EF-Tu, relative to EGDe in mid-log phase in BHI. Data were obtained from three (a-l) and four (m) independent experiments and are presented as mean ± SD (a, k-m) or as median ± interquartile (box) and extreme values (lines) (b-j). Curves were fitted with a Gompertz model and the lag phases (k) for each pair of *Lm* strains were compared with the extra sum-of-squares F test (a). CFU in competition assays are compared with the Wilcoxon matched-pairs signed rank test (b-j) and samples compared with the Kruskal-Wallis test (f, k-m). ns: *p*>0.05, *: *p*<0.05, **: *p*<0.01, ***: *p*<0.001, ****: *p*<0.0001.

**Extended Data Fig. 4.**
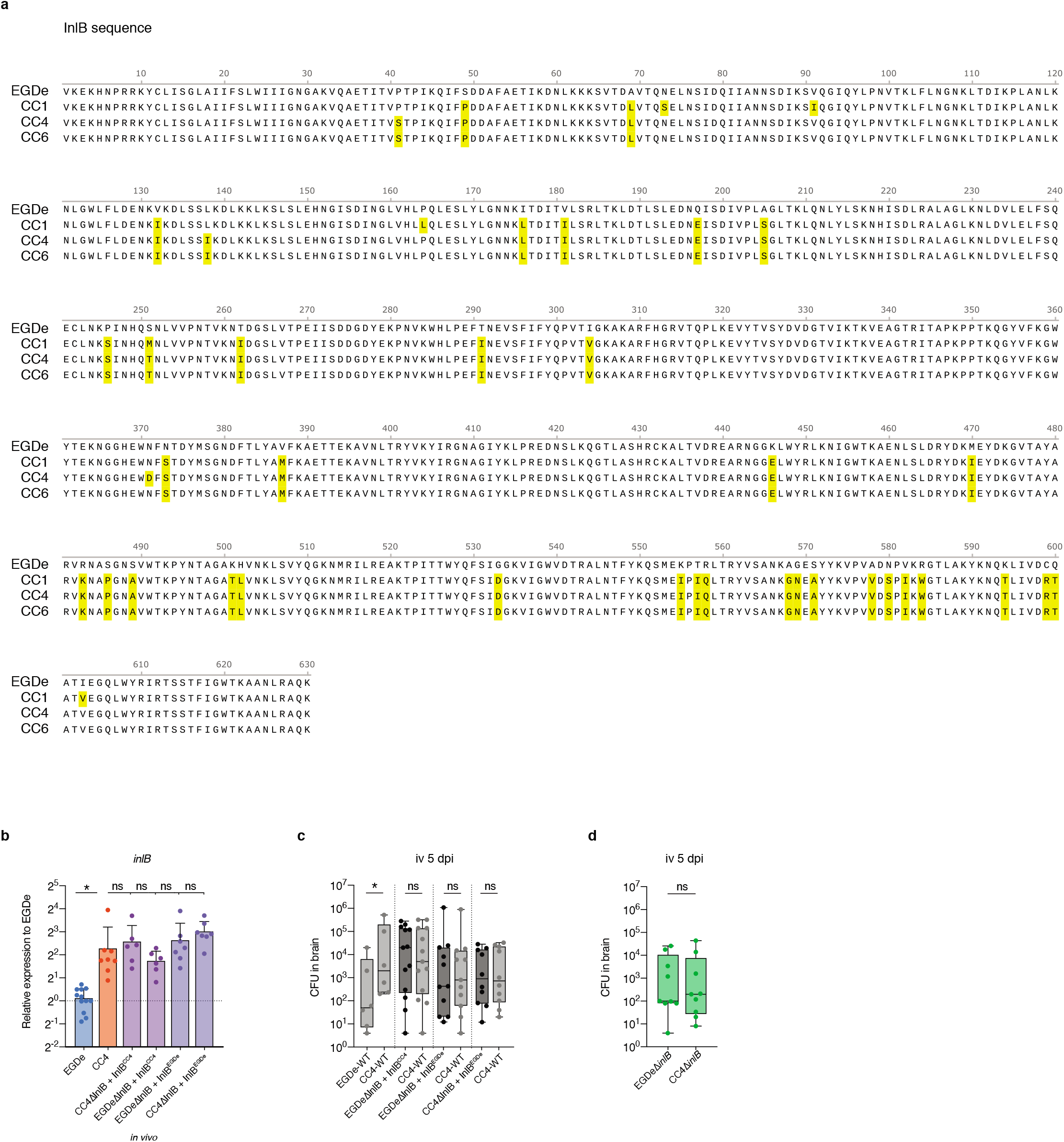
Levels of expression of InlB, and not allelic differences with EGDe, explain enhanced neuroinvasiveness of hypervirulent CC1, CC4 and CC6 strains. **(a)** Protein sequence alignment of InlB alleles from EGDe and from CC1, CC4 and CC6 strains. Mismatches are indicated in yellow. **(b)** Transcription levels of *inlB,* relative to EGDe, in infected splenocytes 2 days after iv inoculation with 2×10^5^ CFU in KIE16P mice of EGDe-WT, CC4-WT and strains complemented with either InlB from EGDe or from CC4. **(c)** Bacterial load in brain of KIE16P mice 5 days after iv inoculation with 10^4^ CFU of a 1:1 mix of the indicated bacterial strains, related to Fig. 2j. **(d)** Bacterial load in brain of KIE16P mice 5 days after iv inoculation with 2×10^4^ CFU of a 1:1 mix of EGDeΔ*inlB* and CC4Δ*inlB*, related to Fig. 2j. Data were obtained from three independent experiments and are presented as mean ± SD (b) or as median ± interquartile (box) and extreme values (lines) (c-d). CFU in competition assays are compared with the Wilcoxon matched-pairs signed rank test (c-d) and samples compared with the Kruskal-Wallis test (b). ns: *p*>0.05, *: *p*<0.05.

**Extended Data Fig. 5.**
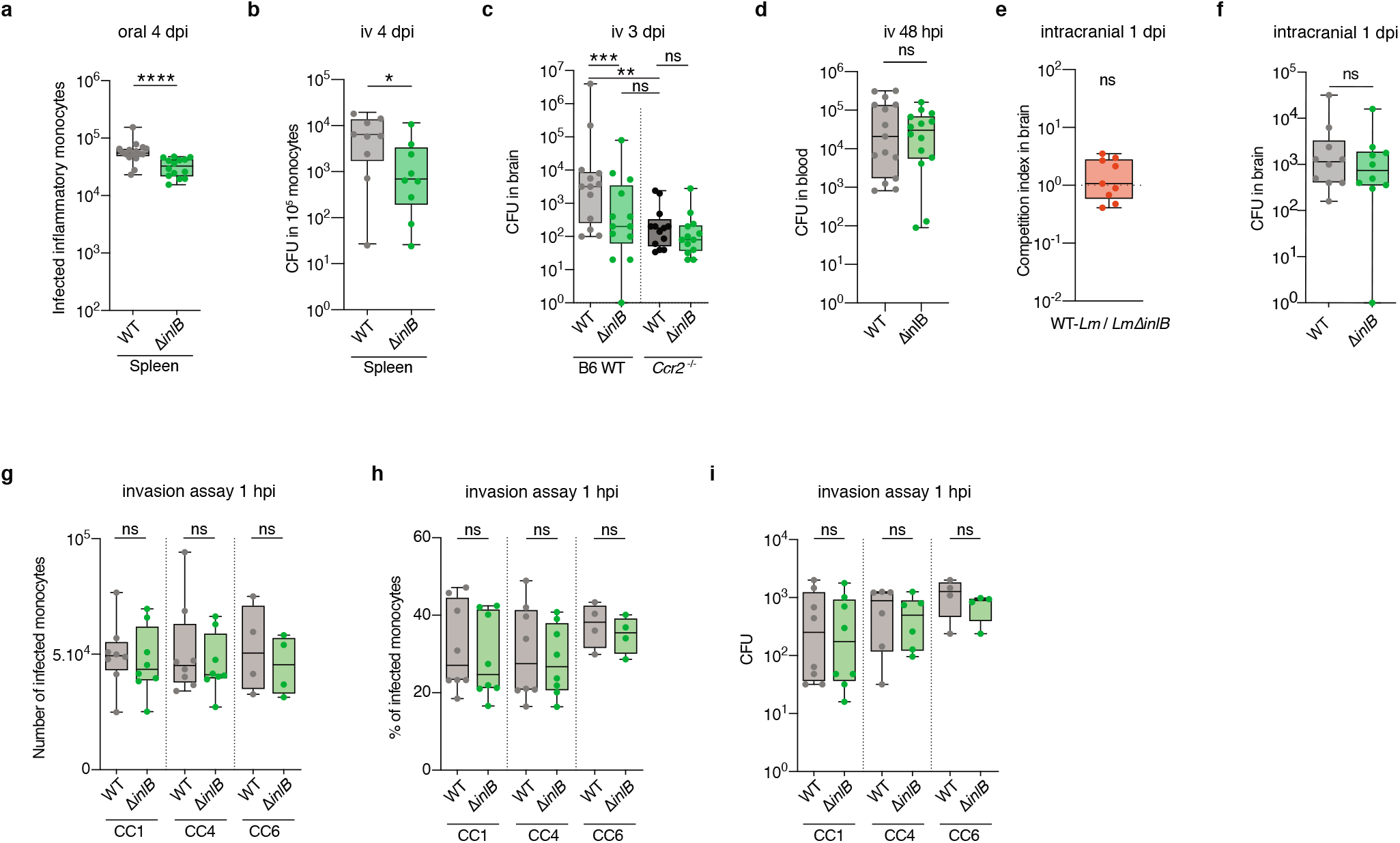
InlB is not involved in *Lm* invasion in monocytes. **(a)** Number of infected monocytes in the spleen of KIE16P mice 4 days after oral inoculation with 2×10^8^ CFU of CC4-WT or CC4Δ*inlB*. **(b)** Bacterial enumeration from sorted monocytes retrieved from KIE16P mice iv infected for 4 days with 10^4^ CFU of CC4-WT or *CC4ΔinlB.* **(c)** Bacterial load in brain of control or *Ccr2^-/-^* mice 3 days after iv inoculation with 10^4^ CFU of 1:1 CC4-WT and *CC4ΔinlB.* **(d)** Bacterial load in the brain 48 hours after iv inoculation of KIE16P mice with 5× 10^5^ CFU of either CC4-WT strain or CC4Δ*inlB.* **(e, f)** Competition index (e) and bacterial load (f) in the brain of KIE16P mice 1 day after intracranial inoculation with 10^2^ CFU of 1:1 mix of CC4-WT and CC4Δ*inlB.* **(g-i)** Number of infected monocytes (g), percentage of infected monocytes (h) and bacterial load (i) in monocytes 1 hour after *in vitro* infection of primary bone marrow mouse monocytes with WT-*Lm* or *ΔinlB* isogenic mutant, at a MOI of 5. Data were obtained from three (d-f) and four (a-c, g-i) independent experiments and are presented as median ± interquartile (box) and extreme values (lines). Samples are compared with the unpaired Mann-Whitney test (a-b, d, g-i) and the Kruskal-Wallis test (c), and CFU in competition assays are compared with the Wilcoxon matched-pairs signed rank test (c, e-f). ns: *p*>0.05.

**Extended Data Figure 6.**
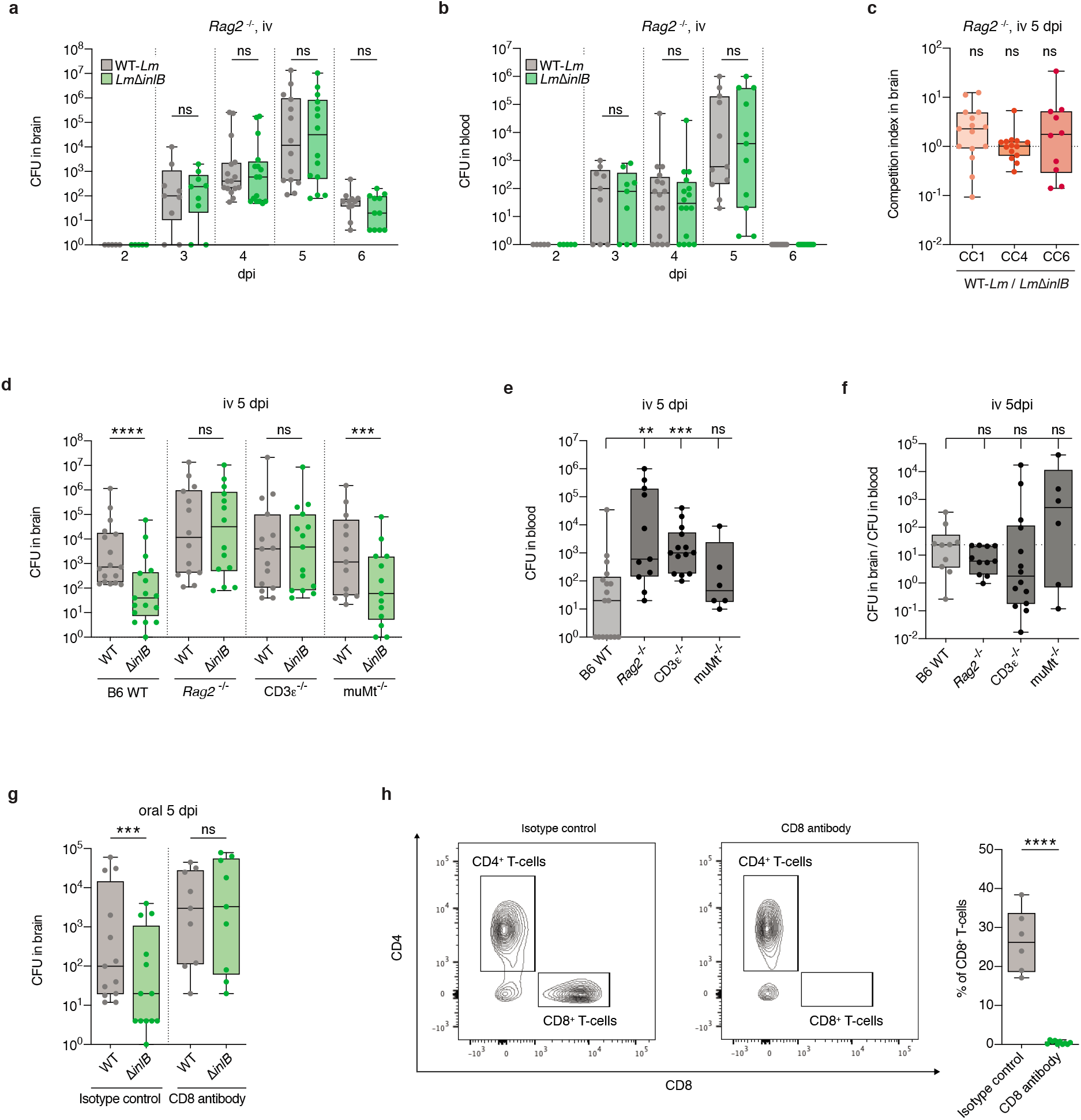
InlB-mediated *Lm* neuroinvasiveness is abrogated in CD8^+^ T-cells deficient mice. **(a, b)** Bacterial load in brain (a) and in blood (b) of *Rag2^-/-^* mice after iv inoculation with 10^4^ CFU of a 1:1 mix of CC4-WT and CC4Δ*inlB*, related to Fig.3f **(c)** Competition index in brain of *Rag2^-/-^* mice 5 days after iv inoculation with 10^4^ CFU of a 1:1 mix WT strain and *ΔinlB* isogenic strains. **(d, e)** Bacterial load in brain (d) and in blood (e) 5 days after iv inoculation with 10^4^ CFU of 1:1 CC4-WT strain and *CC4ΔinlB* isogenic mutant in control mice and in mice lacking functional T (CD3ε^-/-^), B lymphocytes (muMt^-/-^) or both *(Rag2^-/-^),* related to Fig. 3g. **(f)** Ratio of brain/blood bacterial load in control, *Rag2^-/-^* CD3ε^-/-^ and muMt^-/-^ mice, related to Fig. 3g. **(g)** Bacterial load in brain of KIE16P mice 5 days after oral inoculation with 2×10^8^ CFU of 1:1 CC4-WT and *CC4ΔinlB* after CD8^+^ T-cells depletion, related to Fig. 3h. **(h)** Representative dot plots (left) and proportion of CD8^+^ T-cells (right) among CD45^+^ CD3^+^ cells in the spleen, after CD8^+^T-cells depletion, related to Fig. 3h. Data were obtained from three independent experiments and are presented as median ± interquartile (box) and extreme values (lines). CFU in competition assays are compared with the Wilcoxon matched-pairs signed rank test (a-d and g) and samples are compared with the Kruskal-Wallis test (e, f) and with the Mann-Whitney test (h). ns: *p*>0.05, **: *p*<0.01, ***: *p*<0.001, ****: *p*<0.0001.

**Extended Data Figure 7.**
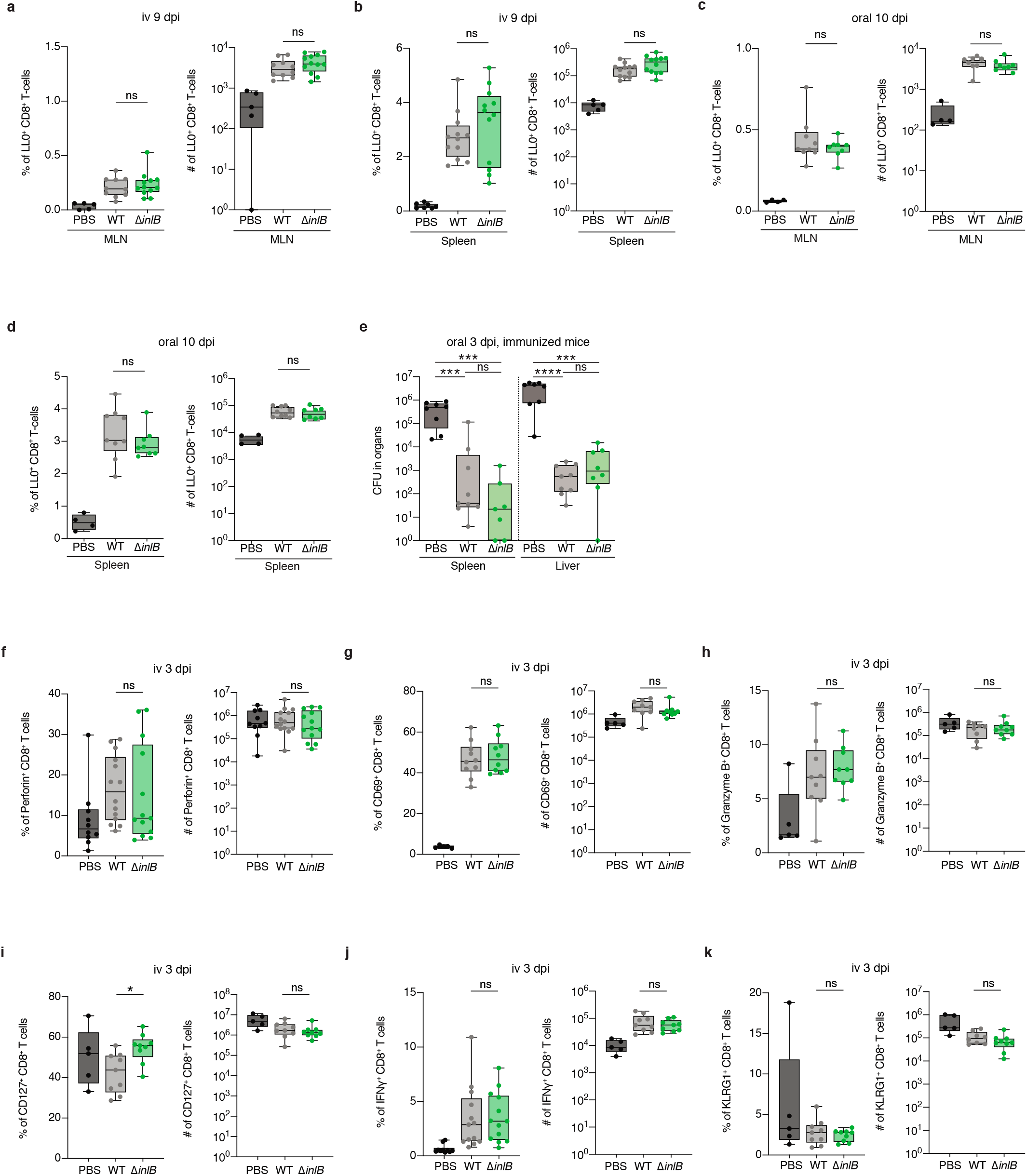
InlB does not alter the induction and differentiation of specific anti-*Lm* CD8^+^ T-cells. **(a, b)** Percentage (left) and number (right) of LLO-specific CD8^+^ T-cells in mesenteric lymph nodes (MLN) (a) and spleen (b) of BALB/c mice 9 days after iv inoculation with 1×10^3^ of CC4-WT strain or CC4*ΔinlB*. **(c, d)** Percentage (left) and number (right) of LLO-specific CD8^+^ T-cells in mesenteric lymph nodes (MLN) (c) and spleen (d) of iFABP-hEcad mice 10 days after oral inoculation with 2×10^7^ of CC4-WT strain or *CC4ΔinlB.* **(e)** Bacterial load in spleen and liver 3 days after oral inoculation with 1×10^9^ CFU of CC4-WT in KIE16P mice challenged 30 days before with 5×10^7^ CFU of CC4-WT strain or *CC4ΔinlB.* **(f-k)** Percentage (left) and number (right) of Perforin^+^ (f), CD69^+^ (g), Granzyme-B^+^ (h), CD127^+^ (i), IFNg^+^ (j) and KLRG1^+^ (k) CD8^+^ T-cells 3 days after iv inoculation of KIE16P mice with 10^4^ CFU of CC4-WT or *CC4ΔinlB.* Data were obtained from three independent experiments and are presented as median ± interquartile (box) and extreme values (lines). Samples are compared with the Mann-Whitney test. ns: *p*>0.05, ***: *p*<0.001, ****: *p*<0.0001.

**Extended Data Figure 8.**
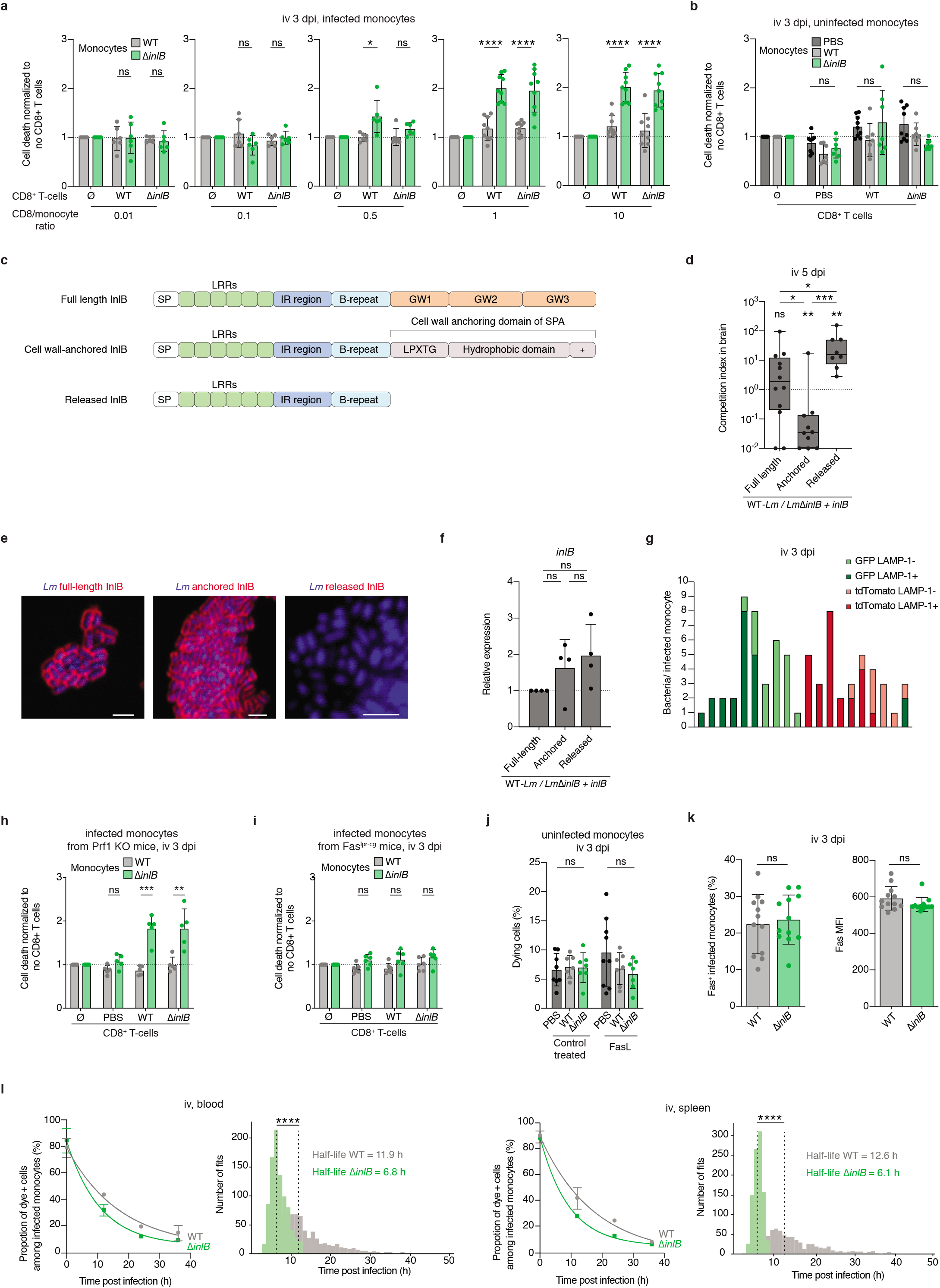
Membrane-associated InlB protects infected monocytes from CD8^+^ T cells-mediated cell death. **(a)**Level of caspase-3 cleavage of infected spleen monocytes, harvested 3 days after iv infection with 10^4^ CFU of CC4-WT or *CC4ΔinlB* of KIE16P mice, and incubated with CD8^+^ T-cells from similarly infected (WT and *ΔinlB)* or control (PBS) mice at the indicated effector to target ratio, related to Fig. 3j. Results are normalized to the level of caspase-3 cleavage in absence of CD8^+^ T cells. (**b**) Level of caspase-3 cleavage of uninfected spleen monocytes, harvested 3 days after iv infection with 10^4^ CFU of CC4-WT or *CC4ΔinlB* of KIE16P mice, and incubated with CD8^+^ T-cells from similarly infected (WT and *ΔinlB*) or control (PBS) mice at an effector to target ratio of 5, related to Fig. 3j. Results are normalized to the level of caspase-3 cleavage in absence of CD8^+^ T cells. (**c**) Schematic representation of WT InlB and its anchored and released variants. (**d**) Competition index in the brain of KIE16P mice 5 days after iv inoculation with 10^4^ CFU of 1:1 CC4-WT and CC4*ΔinlB* transformed with a plasmid expressing either full-length WT InlB, cell wall-anchored InlB or released InlB. (**e**) Representative fluorescence microscopy images of centrifugated CC4*ΔinlB* transformed with a plasmid expressing either full length InlB (left panel), anchored InlB (central panel) or released InlB (right panel). Scale bars: 5 μm. (**f**) Transcription level of *inlB* in *CC4ΔinlB* transformed with a plasmid expressing InlB variants in mid-log phase in BHI, relative to *CC4ΔinlB* expressing full length InlB. (**g**) Proportion of GFP-or tdTomato-expressing bacteria, co-localizing or not with LAMP-1, in 20 infected monocytes harvested 3 days after iv inoculation of KIE16P mice with 10^4^ CFU of 1:1 mix of CC4-WT expressing GFP or tdTomato. (**h, i**) Level of caspase-3 cleavage of infected spleen monocytes, harvested from *Prf1 KO* (h) or *Fas^lpr-cg^* (i) mice, 3 days after iv inoculation with 10^4^CFU of CC4-WT or CC4*ΔinlB*, incubated with CD8^+^ T-cells from similarly infected (WT and *ΔinlB)* or control (PBS) mice, at an effector to target ratio of 5. (**j**) Level of caspase-3 cleavage of non-infected spleen monocytes, harvested from KIE16P mice iv infected for 3 days with 10^4^ CFU of CC4-WT or CC4Δ*inlB*, incubated *ex vivo* with Fas ligand, related to Fig. 3k. (k) Percentage of infected spleen monocytes expressing Fas at their surface (left), and the mean fluorescence intensity (MFI) of Fas signal (right), 3 days after iv inoculation of KIE16P mice with 10^4^ CFU of CC4-WT or CC4*ΔinlB*. (l) Proportion of dye-positive infected monocytes in the blood and the spleen after iv infection of KIE16P mice with 10^4^ CFU of CC4-WT or *CC4-ΔinlB*. The repartition of the estimated half-lives and the median are shown besides each graph. Data were obtained from three independent experiments and are presented as mean ± SD (a, b, f, h-l) or median ± interquartile (box) and extreme values (lines) (d). Samples are compared with an unpaired student *t*-test (a, b, f, h-k), CFU in competition assays are compared with the Wilcoxon matched-pairs signed rank test (d), samples are compared with the Kruskal-Wallis test (d) and distribution of estimated half-lives (i) are compared with a Mood test. ns: *p*>0.05, *: *p*<0.05, ***: *p*<0.001, ****: *p*<0.0001.

**Extended Data Figure 9.**
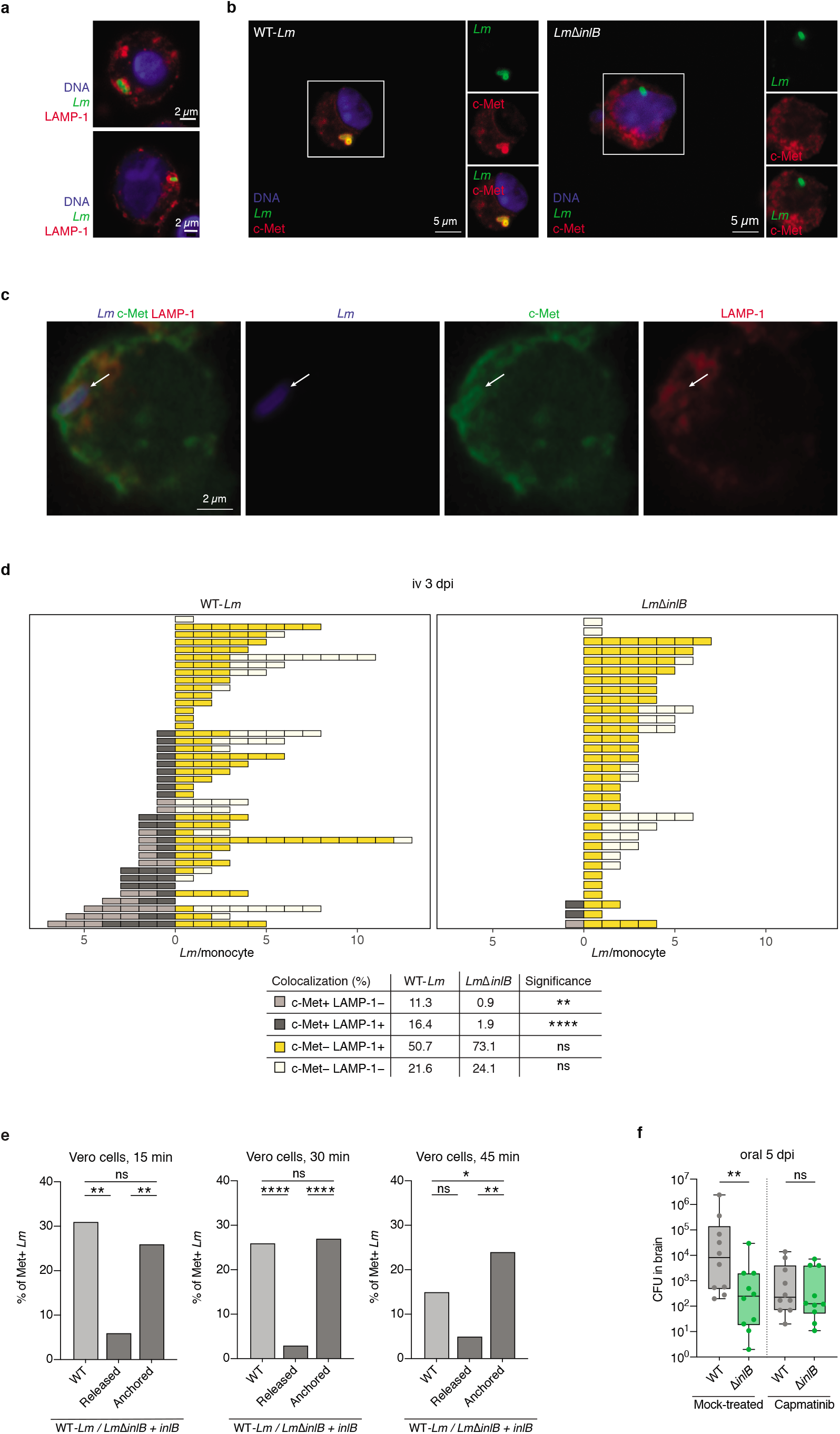
InlB recruits c-Met in infected monocytes. **(a-c)** Representative fluorescence microscopy images of spleen monocytes harvested from KIE16P mice iv infected for 4 days with 10^4^ CFU of CC4-WT or *CC4ΔinlB,* showing intra-vacuolar *Lm* surrounded with LAMP-1 (a), co-localizing with c-Met (b) and co-localizing with both c-Met and LAMP-1 (c). (a and b) are maximum intensity projection over a *z*-stack. (c) is a confocal single plane image. **(d)** Quantification of intracellular *Lm* co-localizing or not with c-Met and LAMP-1 in infected spleen monocytes harvested from KIE16P mice iv infected for 4 days with 10^4^ CFU of CC4-WT or CC4*ΔinlB*. Individual cells are plotted in top panel and samples are compared in bottom panel. **(e)** Percentage of *Lm* co-localizing with c-Met *in vitro* in Vero cells 15 min (left), 30 min (middle) and 45 min (right) after infection at MOI 50 with *CC4ΔinlB* expressing either WT InlB, released InlB or cell wall-anchored InlB. **(f)** Bacterial load in the brain 5 days after oral inoculation with 2×10^8^ CFU of 1:1 of CC4-WT and CC4Δ*inlB*, in KIE16P mice treated with capmatinib, related to Fig. 4a. Data were obtained from three independent experiments (e-f) or from three microscopic field of views (d). Median number of bacteria in each intracellular compartment were compared with the Mann-Whitney test (d), proportions of c-Met associated bacteria (e) with Fischer’s exact test, and CFU in competition assays (f) compared with the Wilcoxon matched-pairs signed rank test. ns: *p*>0.05, *: *p*<0.05, **: *p*<0.01, ****: *p*<0.0001.

**Extended Data Figure 10.**
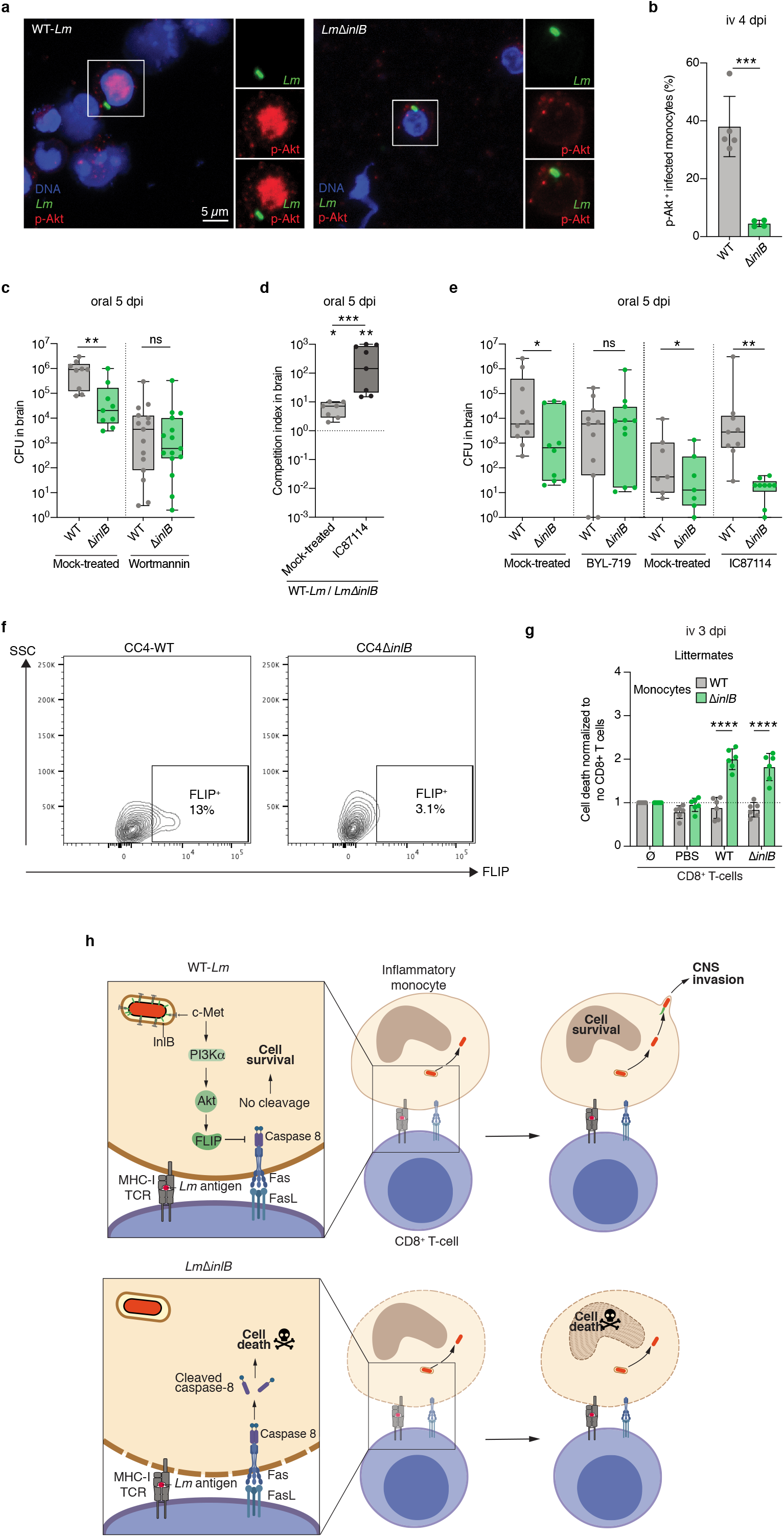
InlB-mediated neuroinvasion involves the c-Met/PI3Kα/FLIP pathway in infected monocytes. **(a)** Representative fluorescence microscopy images of spleen monocytes harvested from KIE16P mice iv infected for 4 days with 10^4^ CFU of CC4-WT or CC4Δ*inlB*, showing phosphorylation of Akt. Images are maximum intensity projection over a *z*-stack. **(b)** Proportion of infected spleen monocytes positive for p-Akt signal 4 days after iv inoculation of KIE16P mice with 10^4^ CFU of CC4-WT or *CC4ΔinlB.* **(c)** Bacterial load in brain 5 days after oral inoculation with 2×10^8^ CFU of 1:1 of CC4-WT and *CC4ΔinlB,* in KIE16P mice treated with wortmannin, related to Fig. 4d. **(d)** Competition index in brain 5 days after oral inoculation with 2×10^8^ CFU of 1:1 of CC4-WT and *CC4ΔinlB,* in KIE16P mice treated with PI3Kδ inhibitor (IC87114). **(e)** Bacterial load in the brain 5 days after oral inoculation with 2×10^8^ CFU of 1:1 of CC4-WT and *CC4ΔinlB,* in mice treated with BYL-719 or IC87114, related to Fig. 4e and Extended Data Fig. 10d. **(f)** Representative dot plot of FLIP expression in infected inflammatory spleen monocytes, 3 days after iv inoculation with 10^4^ CFU of CC4-WT or CC4*ΔinlB*, related to Fig. 4f, g. **(g)** Level of caspase-3 cleavage of infected spleen monocytes, harvested 3 days after iv infection with 10^4^ CFU of CC4-WT or CC4*ΔinlB* of tamoxifen-treated *Rosa26*-CreER^T2^ × *Cflar^+-+^* (FLIP^+/+^) littermate mice and incubated with CD8^+^ T-cells from similarly infected mice at an effector to target ratio of 5, related to Fig. 4h. **(h)** Representation of InlB-activated pathway of infected monocytes survival to Fas-mediated cell death. Data were obtained from three independent experiments and are presented as median ± interquartile (box) and extreme values (lines) (c-e) or mean ± SD (b, g). CFU in competition assays are compared with the Wilcoxon matched-pairs signed rank test (c-e) and samples with a Mann-Whitney test (c-e) or an unpaired student *t*-test (b, g). ns: *p*>0.05, *: *p*<0.05, **: *p*<0.01, ***: *p*<0.001, ****: *p*<0.0001.

**Movie S1. Polymerization of actin comet tail by *Lm* within a monocyte adhering to the blood-brain barrier.**

CX3CR1^GFP/+^ E16P KI humanized mice were infected with 5×10^5^ CFUs of CC6 (isolate 2009-01092) via iv route. Mice were sacrificed 48 hours post infection. CX3CR1^+^ are labeled in green, *L. monocytogenes* in red, actin (phalloidin) in white and nuclei (Hoechst) in blue. Forty-six optical sections of a 20 μm thick brain sample were imaged with a Zeiss LSM700 confocal microscope. 3D reconstruction was performed using the Arivis Vison 4D software.

## Notes

### Competing Interest Statement

The authors have declared no competing interest.

