## Supplementary material for "Bacterial inhibition of CD8^+^ T-cells mediated cell death promotes neuroinvasion and within-host persistence": Methods

### Material and methods

#### Mice

Animal experiments were performed according to the Institut Pasteur guidelines for laboratory animals' husbandry and in compliance with European regulation 2010/63 EU. All procedures were approved by the Animal Ethics Committee of Institut Pasteur, authorized by the French Ministry of Research and registered under #11995-201703115103592 and #14644-2018041116183944.

C57BL/6JRj and BALB/c mice were purchased from Janvier Labs (France) and bred at Institut Pasteur. KIE16P mice expressing humanized E16P E-cadherin[7], iFABP-hEcad mice[15], *Rag2*$^{-/-}$ mice[45], CD3ε$^{-/-}$ mice[46], muMt$^{-/-}$[47], *Ccr2*$^{-/-}$ mice[12], Rosa26CreER$^{T2}$[48] and Rosa26iDTR[49] were bred at Institut Pasteur. *Fas*$^{lpr\text{-}cg}$ carrying a spontaneous mutation at the Fas locus[50], and Prf1 KO mice deleted for the perforin gene[51] were obtained from Frederic Rieux-Laucat and Fernando Sepulveda, Institut Imagine, Paris, respectively. *Cflar*$^{flox/flox}$ (FLIP$^{f/f}$) mice were obtained from Richard M. Pope (Northwestern University, USA)[52], Met$^{flox/flox}$ from Alain Eychene (Institut Curie, Paris)[53] and *LysM*CreER$^{T2}$ from Florian Greten (Georg Speyer Haus, Germany)[54].

All experiments were performed on mice between 7 and 10 weeks of age, randomly assigned to each different condition. Unless stated otherwise in the figure legends, mice refer to KIE16P mice.

#### Bacterial strains

All bacterial strains used in this study are presented in Extended Data Table S. To obtain growth curves, bacteria grown overnight in BHI (with chloramphenicol if needed), at 37°C and 200 rpm, were diluted 1:100 in BHI (with chloramphenicol if needed), incubated at 37°C and 200 rpm, and $OD_{600}$ was measured every hour for 9 hours. Growth curves were fitted using Gompertz model to compare parameters between each strain.

#### Mutagenesis and plasmids

The oligonucleotide primers and plasmids used in this study are listed in Extended Data Table 2 and 3, respectively. Deletion mutants were constructed as described by Monk et *al.*[55]. The flanking regions of the target genes were PCR amplified. After purification, the fragments were stitched together by sequence overlap extension then cloned into the pMAD shuttle vector[56]. The vector was then electroporated into electrocompetent *Lm* cells. After plasmid integration and excision by sequence homology, gene deletion was verified by sequencing the PCR product of the target region.

For *Lm*Δ*inlA* mutants, to avoid alteration of InlB expression in the operon, the *inlA* gene with flanking regions was PCR amplified and cloned into pLR16-pheS plasmid[57] (kind gift from Prof. Anat Herskovits, Tel Aviv University) and a point mutation was introduced via PCR in the *inlA* gene resulting in a premature stop codon. The plasmid was then purified and electroporated into electrocompetent *Lm*Δ*inlA* bacteria. After plasmid integration and excision by sequence homology, allele replacement was verified by sequencing the PCR product of the target region.

The integrative pAD backbone[58] was used to allow expression of TdTomato and GFP. First an EcoRV restriction site was added in pAD between the 5'UTRhly and the ATG start codon, generating the plasmid pCMC12. TdTomato and GFP sequences were codon-optimized for expression in *Lm* using Optimizer (http://genomes.urv.es/OPTIMIZER/), synthesized by Eurofins with EcoRV and SalI flanking restriction sites and inserted in pCMC12 between EcoRV and SalI sites. *inlB* sequences were amplified from EGDe (MBHL0005), CC4 (MBHL0257) and *Listeria innocua* BUG1642 (MBHL0052, expressing a construct where InlB anchoring region has been replaced by the anchoring domain of *S. aureus* protein A[17]) genomic DNAs and all were inserted in pCMC12 between EcoRV and SalI sites. The β-lactamase construct was described in Quereda et *al*[59]. All constructs were electroporated into electrocompetent *Lm* cells. Integration was verified by PCR.

**Infections and bacterial enumerations**

Bacterial inocula were prepared by centrifugation of a bacterial culture grown in BHI, at 37°C and shaken at 200 rpm, until $OD_{600}$ of 0.8 ($8.10^8$ viable bacteria per mL), washed in PBS and resuspended in PBS at the appropriate dilution. For oral inoculation, 0.2 mL of bacteria were mixed to 0.3 mL of PBS containing 50 $mg.mL^{-1}$ of $CaCO_3^-$ (Sigma) and injected intragastrically via a feeding needle (ECIMED) to isoflurane-anesthetized mice. For intravenous infection, 0.1 mL of bacteria were injected in the tail vein using a 25G needle. For intracerebral infections, mice were anesthetized in 3% isoflurane, then received 10 µl of bacterial suspension by intracranial injection using a 26G needle, inserted approximately 2 mm anterior to the bregma, 1 mm laterally and 1.5 mm ventrally.

For immunization experiments, mice were first infected with WT or Δ*inlB* strains by oral gavage with $5.10^7$ CFU. After 30 days, mice were then infected intragastrically for the second challenge with $2.10^9$ WT bacteria for 3 days.

At indicated times, mice were euthanized and their organs aseptically harvested. Intestine, colon and cecum were incubated with gentamicin 100 µg/mL for 2 hours to eliminate extra-tissular bacteria. All organs were homogenized in PBS with a tissue homogenizer (UltraTurrax T-25 basic, IKA works). Serial dilutions of the homogenate were plated onto BHI agar (or ALOA for feces and intestinal content) and CFUs enumerated. CFUs are expressed per mL for blood and per whole organ otherwise.

Competition index experiments were performed as described in Disson *et al*[60]. Briefly, a 1:1 ratio of wild-type bacteria, expressing TdTomato, and mutant bacteria, expressing GFP, was injected into mice, confirmed by CFUs enumeration of the inoculum onto BHI agar. WT and mutant bacteria CFUs were distinguished by colonies' color onto BHI agar. Competition index is calculated as the ratio of WT versus mutant CFUs in each organ. Except stated otherwise, competition indexes were performed 5 days post-inoculation, a time point late enough to have consistent CNS invasion and induction of adaptive immune responses but before humane endpoints were reached. Whenever the mouse strains were permissive to *Lm* oral infection, intragastric inoculation was used for competition index assays.

**Drug treatments of mice**

All drugs used in this study and their mode of delivery are presented in Extended Data Table 4.

**RNA isolation and qRT-PCR**

For *in vitro* analysis, bacteria were grown in BHI at 37°C and 200 rpm until $OD_{600}$ 0.8. They were then centrifuged, lysed in resuspension buffer (10% glucose, 12.5 mM Tris, 10 mM EDTA in nuclease-free water), transferred into Precellys tube (Ozyme) containing 0.1 mm ceramic beads (Ozyme) and acid phenol (Sigma) and homogenized using a Precellys 24 apparatus (Ozyme). Aqueous phase was transferred into a new tube containing TRIzol (Invitrogen) and chloroform (Sigma), shaken and centrifuged at maximum speed at 4°C for 15 min. Aqueous phase was transferred into a new tube containing chloroform, shaken and centrifuged at maximum speed at 4°C for 15 min. Aqueous phase was transferred into a new tube containing 1 volume of isopropanol (Sigma) and 0.1 volume of 3M sodium acetate (Sigma), incubated for at least 20 min at -20°C.

For *in vivo* analysis, spleens were harvested aseptically and single cells suspensions were obtained by homogenizing through a 40 µm cell strainer. Cells were lysed in TRIzol and chloroform, shaken and centrifuged at maximum speed at 4°C for 15 min. Aqueous phase was transferred into a new tube containing 1 volume of isopropanol, incubated for at least 20 min at -20°C.

Both for *in vitro* and *in vivo* samples, RNAs were pelleted at maximum speed at 4°C for at least 20 min and washed three times in 80% ethanol (Sigma), air-dried and dissolved in RNase-free water. Total RNA was reverse-transcribed using hexameric random primers (Invitrogen) and M-MLV reverse transcriptase (Invitrogen) following manufacturer's instructions. Quantification of gene expression was performed using Power SYBR™ Green PCR Master Mix, a Step-One Real-Time PCR apparatus (both from Applied Biosystems) and primers listed in Extended Data Table 2. Expression of *inlA* and *inlB* was normalized to that of *gyrB* and compared to EGDe using the ΔΔCt method.

**Flow cytometry**

Mice were euthanized at the indicated times post-infection. Functional characterization of $CD8^+$ T cells and infected monocytes were performed after iv inoculation, unless stated otherwise, which allows for less intra-animal variation and thus reduces the number of animals needed for experiments. Counting and characterization of infected cells were performed at 4 days post-inoculation (both orally and iv), when bacteremia is the highest. For orally infected mice, the bacteremia being very low even at 4 days post-inoculation, blood of 3 mice was pooled and analyzed as one sample. Spleens, mesenteric lymph nodes (MLN) and blood (by cardiac puncture in heparin-coated syringe) were harvested aseptically. Single cells suspensions were obtained from spleen and MLN by homogenizing through a 40 μm cell strainer. After red blood cells lysis using 1 X RBC lysis buffer (eBiosciences), cells were washed in Cell Staining Buffer (CSB) (BioLegend) before further processing. If required, cells were labeled using the LiveBLAzer™ FRET-B/G Loading Kit with CCF2-AM (Invitrogen), a fluorescent substrate of β-lactamase that can cross the plasma membrane. Presence of β-lactamase-expressing bacteria in cells induces a shift in the fluorescence emission of the CCF2-AM substrate from 518 nM (green) to 447 nM (blue) upon excitation with the 405 nM laser and thus allows for identification of infected cells. Cells were loaded with the CCF2-AM substrate following manufacturer's instructions for 2h30 at room temperature in CSB containing 1mM probenecid (Sigma). After washing, cells were blocked using CD16/32 (BioLegend) for 5 min at room temperature, washed in CSB, stained with the appropriate antibodies (listed in Extended Data Table 5) for 45 min at 4°C and washed in CSB. If no intracellular staining was required, cells were suspended in CSB containing CountCAL beads (Sony) for absolute counting of cells. For intracellular staining, cells were fixed for 20 min at room temperature in IC fixation buffer (eBiosciences), washed three times in 1X permeabilization buffer (eBiosciences), incubated with primary antibodies for 1 hour at room temperature, washed and incubated with secondary antibodies for 45 min at room temperature. After washing, cells were suspended in CSB containing CountCAL beads for absolute counting of cells. Cells were acquired on a Fortessa X-20 SORP apparatus

(BD biosciences) and analyzed using FlowJo software (TreeStar). B-cells were defined as $CD45^{+}$ $CD19^{+}$ $CD3^{-}$ ; $CD8^{+}$ T-cells as $CD45^{+}$ $CD3^{+}$ $CD8^{+}$ $CD19^{-}$ ; LLO-specific $CD8^{+}$ T-cells as $CD45^{+}$ $CD3^{+}$ $CD8^{+}$ LLO-$pentamer^{+}$ $CD19^{-}$ ; $CD4^{+}$ T-cells as $CD45^{+}$ $CD3^{+}$ $CD4^{+}$ $CD19^{-}$ ; granulocytes as $CD45^{+}$ $Ly6G^{+}$ $CD3^{-}$ $CD19^{-}$ ; patrolling monocytes in blood as $CD45^{+}$ $CD11b^{+}$ $CD11c^{+}$ $CD3^{-}$ $CD19^{-}$ $Ly6G^{-}$ $Ly6C^{-}$ ; inflammatory monocytes as $CD45^{+}$ $CD11b^{+}$ $Ly6C^{high}$ $CD3^{-}$ $CD19^{-}$ $Ly6G^{-}$ $CD11c^{-}$ ; macrophages in spleen as $CD45^{+}$ $CD11b^{+}$ $CD11c^{+}$ $CD3^{-}$ $CD19^{-}$ $Ly6G^{-}$ $Ly6C^{-}$ ; dendritic cells as $CD45^{+}$ $CD11c^{+}$ $CD3^{-}$ $CD19^{-}$ $Ly6G^{-}$ $Ly6C^{-}$; infected cells as CCF2-blue and non-infected cells as CFF2-green. Number of cells are expressed per mL of blood and per spleen.

**CTL assays**

Infected mice were euthanized 3 days post-infection, a time point where activated $CD8^{+}$ T cells are already present but prior to cell death induced in infected monocytes. Spleens were harvested aseptically and cells prepared as described above for flow cytometry, resuspended in CSB and sorted on a FACSAria III apparatus (BD Biosciences) into fetal calf serum-containing tubes. Activated $CD8^{+}$ T-cells were defined as $CD45^{+}$ $CD3^{+}$ $CD8^{+}$ $CD69^{+}$ $CD19^{-}$, infected inflammatory monocytes as $CD45^{+}$ $CD11b^{+}$ $Ly6C^{high}$ $CD3^{-}$ $CD19^{-}$ $Ly6G^{-}$ $CD11c^{-}$ CCF2-blue and non-infected inflammatory monocytes as $CD45^{+}$ $CD11b^{+}$ $Ly\text{-}C^{high}$ $CD3^{-}$ $CD19^{-}$ $Ly6G^{-}$ $CD11c^{-}$ CCF2-green. For mock-treated mice, $CD8^{+}$ T-cells and inflammatory monocytes were sorted. After sorting, cells were washed and resuspended in RPMI medium (Invitrogen) containing 10% fetal calf serum. Activated $CD8^{+}$ T-cells and monocytes (infected and non-infected), isolated from independent mice, were co-incubated at the indicated ratio for 80 minutes at 37°C, washed and fixed with IC fixation buffer overnight at 4°C. After three washes in 1X permeabilization buffer, cells were stained with anti-cleaved caspase-3 antibody for 1 hour at room temperature, washed and stained with secondary antibody for 45 minutes at room temperature. After washing, cells were acquired on a X-20 Fortessa SORP apparatus and percentage of monocytes positive for cleaved caspase-3 signal analyzed using FlowJo software.

**Half-life of infected monocytes**

Mice were infected intravenously with $10^4$ *Lm* CFUs. At day 2 post inoculation, single cells suspensions were obtained from spleens of non-infected mice as described in the flow cytometry section. Cells were then labelled with Vybrant DiD solution (Invitrogen™) for 20 minutes at 37°C, according to the manufacturer's instructions. After 3 washes in DMEM-F12 medium, cells were resuspended in PBS and immediately transferred into infected mice intravenously.

Mice were then euthanized at indicated time points post-infection and spleens and blood samples were harvested aseptically. Single cells suspensions were then prepared and loaded with the CCF2-AM substrate as described above. After washing, cells were acquired on a X-20 Fortessa SORP apparatus and the percentage of infected monocytes positive for Vybrant DiD signal was analyzed using FlowJo software.

**Estimation of half-life times from exponential fits**

Means and standard deviations of infected monocytes were estimated at each time point (0, 12, 24, 36 hour) assuming they are drawn from a normal distribution. At each time point, random values calculated from the normal distribution were drawn and a new decay curve was simulated. To estimate the half-life of the infected monocytes, 1000 curves were simulated and fitted with an exponential decay. Non-convergent fits or those with $R^2 < 0.97$ were discarded; half-life times were calculated from the remaining fits, and distributions are inferred from the [0 ; 95] percentiles. The differences between half-life were assessed with a non-parametric Mood test in which the null hypothesis assumes that the compared samples come from populations with the same median.

**Fas ligand treatment**

Infected mice were euthanized 3 days post-infection and spleens were harvested aseptically. Cells were prepared and sorted in the same conditions as for CTL assays. Infected and non-infected inflammatory monocytes were sorted into fetal calf serum-containing tubes. After sorting, cells were washed and resuspended in RPMI medium containing 10% fetal calf serum. Cells were then treated with either HA antibody as control (Cell Signaling) or with HA antibody plus recombinant mouse Fas ligand/TNFSF6 (R&D systems) for 80 minutes at 37°C. After washing, cells were fixed overnight in IC fixation buffer. After three washes in 1X permeabilization buffer, cells were stained with anti-cleaved caspase-3 antibody for 1 hour at room temperature, washed and stained with secondary antibody for 45 minutes at room temperature. After washing, cells were acquired on a X-20 Fortessa SORP apparatus and percentage of monocytes positive for cleaved caspase-3 signal analyzed using FlowJo software.

**Caspase 8 activity assay**

Mice were inoculated intravenously, treated with either BYL-719 or Capmatinib and euthanized at 3 days post-infection. Single cells suspensions were prepared from spleen as described in the flow cytometry section. After CCF2-AM loading, blocking with CD16/32 and staining with the appropriate antibodies, cells were incubated for 30 min at 37°C with Red-IETD-FMK from CaspGLOW™ Red Active Caspase-8 Staining Kit (Clinisciences), washed and suspended in Wash buffer containing CountCAL beads for absolute counting of cells. Cells were immediately acquired on a Fortessa X-20 SORP apparatus and analyzed using FlowJo software.

**Transfer of infected monocytes**

Infected KIE16P mice were euthanized 3 days post-infection and spleens were harvested aseptically. Cells were prepared as described above and sorted in the same conditions as for CTL assays. Infected monocytes from 6 mice were sorted into fetal calf serum-containing tubes, pooled, washed and injected into a naïve mouse treated with gentamicin, corresponding to an inoculum of $10^4$ live CFUs. Two days post-injection, the mouse was euthanized, organs harvested aseptically and bacterial enumerated as described above.

*LysM*-CreER$^{T2+/-}$ × *Rosa26*-iDTR$^{+/-}$ and their littermates were infected intravenously with CC4-*Lm*, treated with tamoxifen to allow for Cre expression, euthanized 4 days after infection and spleens harvested aseptically. Cells were prepared as described above and sorted in the same conditions as for CTL assays. Infected monocytes from 3 mice were sorted into fetal calf-serum containing tubes, pooled (corresponding to an inoculum of $2x10^4$ live CFUs), washed and injected into a naïve recipient mouse (same genotype as donor mice) treated with both diphtheria toxin and gentamicin, to kill Cre-expressing monocytes and extracellular bacteria. Four days post-injection, mice were euthanized, organs harvested aseptically and bacteria enumerated as described above.

**Bacterial enumeration in infected monocytes**

Infected mice were euthanized 4 days post-infection, when the bacteremia is the highest, and spleens and blood were harvested aseptically. Cells were prepared as described above and sorted in the same conditions as described for CTL assays. Monocytes were collected into fetal calf serum-containing tubes, washed and resuspended into 0.1% triton, serially diluted in PBS and plated on BHI agar.

**Immunofluorescence labelling for microscopy**

For brain sections microscopy, mice were infected with a higher inoculum, $5.10^5$ CFUs, and euthanized 2 days post-iv inoculation, a time point with enough crossing events to allow analysis but without damages to the blood-brain barrier. Brain hemispheres were fixed in 4% paraformaldehyde in PBS overnight at 4°C then washed in PBS, embedded in 4% agarose and sectioned into 40 µm-thick slices using a vibratome (ThermoScientific, HM 650V). Slices were washed in PBS then incubated for 2 hours in blocking-permeabilization solution (10% goat serum, 4% fetal calf serum and 0.4% Triton X-100 in PBS). Tissues were then labeled with the appropriate primary antibodies (listed in Table S5) overnight at 4°C in mild blocking conditions (4% goat serum, 4% fetal calf serum and 0.4% Triton X100 in PBS), washed in PBS, then incubated with secondary antibodies (listed in Table S5), Hoechst-3342 and Phalloidin-Alexa 647 for 2 hours at room temperature. Tissues were washed in PBS and then mounted on glass slides under coverslips in mounting medium (Invitrogen). The slides are let in obscurity overnight before observation under a Zeiss LM710 or LM700 microscope and acquisition with the ZEN software.

For monocytes microscopy, sorted monocytes were seeded on poly-D-lysine-coated 96 wells-plates, fixed in 4% paraformaldehyde in PBS overnight at 4°C and then washed in PBS. Cells were then permeabilized in 0.1% Triton X-100 for 10 minutes, incubated with blocking solution (5% BSA in PBS) for 30 minutes at room temperature, then labeled with anti-*Lm* antibody for 1 hour at room temperature in PBS-BSA, washed in PBS, and then incubated with anti-rabbit secondary antibody, Hoechst-3342 (ThermoFisher) and Phalloidin-Alexa 647 (ThermoFisher) for 1 hour at room temperature. For c-Met staining, cells were incubated with anti-c-Met in mild blocking conditions (2.5% BSA, 0.2% Triton X-100 in PBS) overnight at 4°C, washed in PBS and incubated with anti-*Lm* overnight at 4°C. For p-Akt and LAMP-1 staining, cells were labelled with primary antibodies solution (anti p-Akt or anti LAMP-1 and anti-*Lm*) in mild blocking conditions (2.5% BSA, 0.2% Triton X-100 in PBS) overnight at 4°C then washed in PBS. Cells were then incubated with appropriate secondary antibodies solution (see Extended Data Table 5) and Hoechst-3342 for 2h at room temperature, washed in PBS and left in PBS at 4°C before observation under a Zeiss LM710 microscope and acquisition with the ZEN software. For microscopy of BHI grown-bacteria, 50 µL of the overnight culture were diluted in 1 mL PBS, spun down in a microfuge for 2 minutes, fixed in 4% paraformaldehyde in PBS for 15 minutes at room temperature then washed in PBS. Bacteria were then permeabilized using 0.5% Triton X100 for 10 min, washed in PBS, then incubated with blocking solution (1% BSA in PBS) for 30 minutes. Next, bacteria were labelled with anti-InlB primary antibody in blocking solution (1% BSA in PBS) for 1 hour at room temperature, washed 3 times in PBS, then incubated with anti-rabbit secondary antibody for 1h at room temperature. After 3 washes in PBS, bacteria were resuspended in Hoechst solution (dilution 1/5000 in PBS) for 15 minutes at room temperature, washed twice with PBS then resuspended in 4 µL PBS. Finally, the bacterial suspension was loaded onto a glass slide coated with 1% agarose gel and a coverslip.

**Immunoblotting**

To assess InlB expression level, bacterial cultures at $OD_{600}$ of 0.8 were centrifuged at 3000 g for 10 minutes. Pellets were then incubated with B-PER™ Complete Bacterial Protein Extraction Reagent (ThermoFisher Scientific) for 15 minutes at room temperature and centrifuged at 16,000 g for 20 minutes to obtain lysates.

Lysates were mixed with reducing sample buffer (NuPAGE, Invitrogen) for electrophoresis and subsequently transferred onto a nitrocellulose membrane. Membranes were then blocked with 5% non fat milk diluted in PBS Tween 0.1 % for 1 hour and incubated with primary antibodies in blocking solution for 2 hours at room temperature. After 1 hour of secondary antibody incubation, immunodetection was performed by using a chemiluminescence kit (Amersham™ ECL™ Prime, GE Healthcare), and bands were revealed using the PXi imaging system (SYNGENE).

***In vitro* monocytes invasion assays**

C57BL/6JRj mice were euthanized and bone marrow was collected aseptically. Cells were washed in PBS, red blood cells lysed as described in the flow cytometry section and monocytes isolated using the mouse Monocyte Isolation Kit (Miltenyi Biotec) following manufacturer's instructions. Cells were incubated overnight at 37°C in RPMI + 10% fetal calf serum and penicillin/streptomycin, washed in RPMI, plated in 96 wells-plate, infected with GFP-expressing *Lm* at MOI 5 for 1 hour at 37°C and treated with 50 µg/mL gentamicin for 1 hour at 37°C. For bacterial enumeration, cells were washed in PBS, lysed in 0.1% triton, serially diluted in PBS and plated onto BHI agar. For flow cytometry analysis, cells were washed, fixed in IC fixation buffer in presence of CountCAL beads, acquired on a X-20 Fortessa SORP apparatus and analyzed using FlowJo software.

***In vitro* Vero cells infection**

Vero cells were seeded on poly-D-lysine-coated 96 wells-plates in DMEM (Invitrogen) + 5% fetal calf serum and penicillin/streptomycin 24 hours prior to infection. On the day of infection, cells were washed three times in DMEM + 0.2% fetal calf serum and incubated in this medium for four hours. Bacteria grown in BHI at 37° and 200 rpm until

$OD_{600}$ 0.8 were centrifuged, washed in PBS, suspended in DMEM and added to the cells at a MOI of 50. After 1 min centrifugation at 200 g, cells were incubated at 37° for the indicated times, fixed for 15 min in 4% PFA and washed three times in PBS. Staining for microscopy was performed as for sorted monocytes in the above section.

**Statistical analysis**

Statistical details and number of replicates are found in the corresponding Figure legends. All statistical tests were two-sided. Analyses were performed using GraphPad Prism 8 Software.

**Extended references**

**Table S1**

| Strains | CLIP number | Collection number | Plasmid | Antibiotic resistance | Comment | Reference |
|---|---|---|---|---|---|---|
| **EGDe** | | MBHL0005 | | | | *(6)* |
| **10403S** | | MBHL0308 | | | | (6) |
| **CC1** | 2007/00596 | MBHL 0237 | | | | (6) |
| **CC4** | 2009/00558 | MBHL 0257 | | | | (6) |
| **CC6** | 2009/01092 | MBHL 0255 | | | | (6) |
| **EGDe β-lactamase** | | MBHL396 | pCMC11 | Chloramphenicol | | This study |
| **CC1 β-lactamase** | 2007/00596 | MBHL398 | pCMC11 | Chloramphenicol | | This study |
| **CC4 β-lactamase** | 2009/00558 | MBHL400 | pCMC11 | Chloramphenicol | | This study |
| **CC6 β-lactamase** | 2009/01092 | MBHL425 | pCMC11 | Chloramphenicol | | This study |
| **EGDe TdTomato** | | MBHL418 | pCMC34 | Chloramphenicol | | This study |
| **CC1 TdTomato** | 2007/00596 | MBHL416 | pCMC34 | Chloramphenicol | | This study |
| **CC4 TdTomato** | 2009/00558 | MBHL412 | pCMC34 | Chloramphenicol | | This study |
| **CC6 TdTomato** | 2009/01092 | MBHL420 | pCMC34 | Chloramphenicol | | This study |
| **EGDe ΔInlB** | | MBHL 0489 | | | | This study |
| **CC1 ΔInlB** | 2007/00596 | MBHL 0490 | | | | This study |
| **CC4 ΔInlB** | 2009/00558 | MBHL 0491 | | | | This study |
| **CC6 ΔInlB** | 2009/01092 | MBHL 0492 | | | | This study |
| **CC1 ΔInlA** | 2007/00596 | MBHL 0493 | | | | This study |
| **CC4 ΔInlA** | 2009/00558 | MBHL 0494 | | | | This study |
| **CC6 ΔInlA** | 2009/01092 | MBHL 0495 | | | | This study |
| **CC4 ΔInlB β-lactamase** | 2009/00558 | MBHL439 | pCMC11 | Chloramphenicol | | This study |
| **CC4 ΔInlB TdTomato** | 2009/00558 | MBHL475 | pCMC34 | Chloramphenicol | | This study |
| **EGDe ΔInlB GFP** | | MBHL449 | pCMC44 | Chloramphenicol | | This study |
| **CC1 ΔInlB GFP** | 2007/00596 | MBHL451 | pCMC44 | Chloramphenicol | | This study |
| **CC4 ΔInlB GFP** | 2009/00558 | MBHL453 | pCMC44 | Chloramphenicol | | This study |
| **CC6 ΔInlB GFP** | 2009/01092 | MBHL455 | pCMC44 | Chloramphenicol | | This study |
| **EGDe ΔInlB + InlB full length from CC4** | | MBHL457 | pCMC58 | Chloramphenicol | | This study |
| **CC4 ΔInlB + InlB full length from CC4** | 2009/00558 | MBHL469 | pCMC58 | Chloramphenicol | | This study |
| **CC4 ΔInlB + released InlB from CC4** | 2009/00558 | MBHL471 | pCMC59 | Chloramphenicol | | This study |
| **CC4 ΔInlB + anchored InlB from CC4** | 2009/00558 | MBHL473 | pCMC60 | Chloramphenicol | | This study |
| **EGDe ΔInlB + InlB full length from EGDe** | | MBHL487 | pCMC61 | Chloramphenicol | | This study |
| **CC4 ΔInlB + InlB full length from EGDe** | 2009/00558 | MBHL488 | pCMC61 | Chloramphenicol | | This study |
| **CC4 ΔActA β-lactamase** | 2009/00558 | MBHL0497 | pCMC11 | Chloramphenicol | | This study |
| **Listeria Innocua InlB anchored** | | MBHL 0052 (BUG1642) | p1B1 | | | (*17*) |
| **CC4 ΔInlAΔInlB** | 2009/00558 | MBHL0496 | | | | This study |
| **CC1** | 2015/00085 | | | | Clinical strain | This study |
| **CC1** | 2016/01406 | | | | Clinical strain | This study |
| **CC1** | 2016/01398 | | | | Clinical strain | This study |
| **CC1** | 2015/00918 | | | | Clinical strain | This study |
| **CC1** | 2016/00717 | | | | Clinical strain | This study |
| **CC1** | 2011/00412 | | | | Clinical strain | This study |
| **CC1** | 2016/01677 | | | | Clinical strain | This study |
| **CC1** | 2015/01910 | | | | Clinical strain | This study |
| **CC1** | 2005/00008 | | | | Clinical strain | This study |
| **CC4** | 2015/00091 | | | | Clinical strain | This study |
| **CC4** | 2015/00895 | | | | Clinical strain | This study |
| **CC4** | 2016/01178 | | | | Clinical strain | This study |
| **CC4** | 2016/01062 | | | | Clinical strain | This study |
| **CC4** | 2015/01543 | | | | Clinical strain | This study |
| **CC4** | 2005/00190 | | | | Clinical strain | This study |
| **CC4** | 2009/00994 | MBHL 0353 | | | Clinical strain | This study |
| **CC4** | 2012/01728 | MBHL 0352 | | | Clinical strain | This study |
| **CC4** | 2013/00255 | MBHL 0350 | | | Clinical strain | This study |
| **CC4** | 2013/00344 | MBHL 0354 | | | Clinical strain | This study |
| **CC6** | 2016/00566 | | | | Clinical strain | This study |
| **CC6** | 2016/00074 | | | | Clinical strain | This study |
| **CC6** | 2016/00783 | | | | Clinical strain | This study |
| **CC6** | 2015/01823 | | | | Clinical strain | This study |
| **CC6** | 2015/00880 | | | | Clinical strain | This study |
| **CC6** | 2015/00001 | | | | Clinical strain | This study |
| **CC6** | 2016/00551 | | | | Clinical strain | This study |
| **CC6** | 2016/01114 | | | | Clinical strain | This study |
| **CC6** | 2015/01823 | | | | Clinical strain | This study |

**Table S2**

| Oligos | sequence | Comment |
|---|---|---|
| CMC15 | GGTTAAAAAATGTAGAAGGAGAGTGATATCCATGAGTAAAGGAGAAGAACTTTTCACTGG | Addition of EcoRV site in pAD to obtain pCMC12-*Forward* |
| CMC16 | CCAGTGAAAAGTTCTTCTCCTTTACTCATGGATATCACTCTCCTTCTACATTTTTTAACC | Addition of EcoRV site in pAD to obtain pCMC12-*Reverse* |
| CMC37 | GCGCCGATATCCATGGTTTCTAAAGGTGAAGAAGTTATTAAAGAATTC | TdTomato (codon-optimized) cloning in pCMC12-*Forward* |
| CMC38 | GCGCCGTCGACTTATTTATATAATTCATCCATACCATATAAGAATAAATGATGAC | TdTomato (codon optimized) cloning in pCMC12-*Reverse* |
| CMC62 | GCGCCGATATCCATGTCTAAAGGTGAAGAATTATTC | GFP (codon-optimized) cloning in pCMC12-*Forward* |
| CMC63 | GCGCCGTCGACTTATTTATATAATTCATCCATACC | GFP (codon-optimized) cloning in pCMC12-*Reverse* |
| CMC195 | GCGCGGATATCCATGAAAGAAAAGCACAACCC | InlB cloning in pCMC12-*Forward* |
| CMC196 | GCGCGGTCGACTTATTTCTGTGCCCTTAAATTAG | InlB cloning in pCMC12-*Reverse* |
| CMC197 | GCGCGGTCGACTTACGTCCCTGCTTCTACTTTTG | InlB without anchoring region cloning in pCMC12-*Reverse* |
| CMC198 | GCGCGGTCGACTTATAGTTCGCGACGACG | InlB with anchoring region of spA cloning in pCMC12-*Reverse* |
| inlA1F | ACGTGTCGACAGATAACACAATCACACCGTGT | inlA1: upstream flanking region of inlA |
| inlA1R | TACACTACTTTATATACACTCCTTTTTCAATAGTTAGAAACA | |
| inlA2F | AGTGTATATAAAGTAGTGTAAAGAGCTAGATGTGGT | inlA2: downstream flanking region of inlA |
| inlA2R | ACGTGGATCCTTTTCAAGTTTGCTAAGGGCTTT | |
| inlB1F | ACGTGTCGACACAACGACTCAAGCAGTAGACT | inlB1: upstream flanking region of inlB |
| inlB1R | TTTTCGTAGGACTATCCTCTCCTTGATTCTAGTTAT | |
| inlB2F | AGAGGATAGTCCTACGAAAAGCTATTTCTAAA | inlB2: downstream flanking region of inlB |
| inlB2R | ACGTGGATCCATGCTATCCACATTTTGGCT | |
| inlAB2R | TTTTCGTAGGTATATACACTCCTTTTTCAATAGTTAGAAACA | inlA1pB2: upstream flanking region of InlA containing overlap with inlB2 |
| inlAB2F | AGTGTATATACCTACGAAAAGCTATTTCTAAA | inlB2pA1: downstream flanking region of InlB containing overlap with inlA1 |
| actA1F | GGGGTCGACTAAAGGTCCACGTCACACCG | actA1: upstream flanking region of actA |
| actA1R | AACCGCTCCTACTACCATCATCGCACGCAT | |
| actA2F | TGATGGTAGTAGGAGCGGTTATCAAAATCATTCA | actA2: downstream flanking region of actA |
| actA2R | TGATGGTAGTAGGAGCGGTTATCAAAATCATTCA | |
| gyrB_F | AAGTATCTGGCGGACTTCACG | qPCR primer pair for gyrB |
| gyrB_R | TCACCACGTTCAAAGCGTTG | |
| inlA_F | CGAAAAATCCTGTGGCACCA | qPCR primer pair for inlA |
| inlA_R | TTTGCGGAAGGTGGTGTAGT | |
| inlB_F | AAGCACAACCCAAGAAGGAA | qPCR primer pair for inlB |
| inlB_R | CGGTGATAGTCTCCGCTTGT | |

**Table S3**

| Plasmids | Comment | Antibiotic resistance | Reference |
|---|---|---|---|
| pAD | chromosomally integrative plasmid | Chloramphenicol | (*58*) |
| pCMC11 | pAD encoding for codon-optimized beta-lactamase anchored to Listeria cell-wall | Chloramphenicol | (*59*) |
| pCMC12 | pAD modified to add an EcoRV restriction site between UTRhly and starting codon | Chloramphenicol | this study |
| pCMC34 | pCMC12 encoding for codon-optimized TdTomato | Chloramphenicol | this study |
| pCMC44 | pCMC12 encoding for codon-optimized GFP | Chloramphenicol | this study |
| pCMC58 | pCMC12 encoding for full length InlB from CC4 | Chloramphenicol | this study |
| pCMC59 | pCMC12 encoding for InlB from CC4 truncated of its anchoring region | Chloramphenicol | this study |
| pCMC60 | pCMC12 encoding for InlB from CC4 with the anchoring region of SpA | Chloramphenicol | this study |
| pCMC61 | pCMC12 encoding for full length InlB from EGDe | Chloramphenicol | this study |
| pMAD | | Erythromicin | (*56*) |
| pMAD-delta-inlA | pMAD+inlA1+inlA2, inlA deletion | Erythromicin | this study |
| pMAD-delta-inlB | pMAD+inlB1+inlB2, inlB deletion | Erythromicin | this study |
| pMAD-delta-inlAB | pMAD+inlA1+inlB2, inlAB deletion | Erythromicin | this study |
| pMAD-delta-actA | pMAD+actA1+actA2, actA deletion | Erythromicin | this study |
| pLR16 | | Chloramphenicol | (*57*) |
| pLR16-inlA-C1474T | introduce premature stop codon in inlA | Chloramphenicol | this study |

**Table S4**

| Drug | Final concentration | Delivery mode | Frequence of delivery | Vehicule | Reference | Supplier | Comments |
|---|---|---|---|---|---|---|---|
| Ciclosporin A | 40 mg/kg | Intraperitoneal | Every day, starting the day prior to infection | Olive oil | Sandimmun | Novartis Pharma | |
| Anti-CD8 antibody | 0.3 mg/mice | Intraperitoneal | 3 days and 1 day prior to infection, day of infection and day 4 post-infection | PBS | BX-BE0061 | Euromedex | Clone 2.43 |
| Isotype control | 0.3 mg/mice | Intraperitoneal | 3 days and 1 day prior to infection, day of infection and day 4 post-infection | PBS | BX-BE0090 | Euromedex | Clone LTF-2 |
| Z-IETD-FMK | 5 mg/kg | Intraperitoneal | Every 2 days, starting the day of infection | DMSO/PBS (1/20) | M3136 | Euromedex | Caspase 8 inhibitor |
| Capmatinib | 3 mg/kg | Oral | Every day, starting the day of infection | DMSO/PEG-400 (1/9) | HY-13404 | CliniSciences | c-Met inhibitor |
| Wortmannin | 1 mg/kg | Intraperitoneal | Every day, starting the day of infection | DMSO/PBS (1/4) | HY-10197 | CliniSciences | Pan PI3K inhibitor |
| BYL-719 | 50 mg/kg | Oral | Every day, starting the day of infection | PEG-400 | HY-15244 | CliniSciences | PI3Ka inhibitor |
| IC87114 | 30 mg/kg | Oral | Twice every day, starting the day of infection | PEG-400 | HY-10110 | CliniSciences | PI3Kd inhibitor |
| Gentamicin | 30 mg/kg | Intraperitoneal or intravenous | Every day, starting the day after infection | PBS | G1272 | Sigma | |
| Tamoxifen | 1 mg/kg | intraperitoneal | Every day, staring 5 days prior to infection | Corn oil | T5648 | Sigma | |
| Diphteria toxin | 25ng/g | Intraperitoneal and intravenous | Intravenous morning of transfer and then IP at time of transfer and 2 days later | PBS | D0564 | Sigma | |

**Table S5**

| Antibody target | Fluorophore | Reference | Supplier | Use |
|---|---|---|---|---|
| CD45 | BV605 | BLE103139 | Ozyme | Flow cytometry |
| CD3 | BV711 or FITC | BLE100241 or BLE100204 | Ozyme | Flow cytometry |
| CD19 | PE-Cy5 | BLE115510 | Ozyme | Flow cytometry |
| CD11b | BUV737 | 564443 | BD Biosciences | Flow cytometry |
| CD11c | BV785 | BLE117336 | Ozyme | Flow cytometry |
| LY-6G | PE-Cy7 or APC-Cy7 | BLE127618 or BLE 127624 | Ozyme | Flow cytometry |
| LY-6C | PE or APC-Cy7 | BLE128008 or BLE128026 | Ozyme | Flow cytometry |
| CD8a | BUV395 or PE-Dazzle594 | 563786 or BLE100762 | BD Biosciences or Ozyme | Flow cytometry |
| CD69 | PE-Cy7 | BLE104512 | Ozyme | Flow cytometry |
| CD4 | PerCP-Cy5-5 | BLE100539 | Ozyme | Flow cytometry |
| CD350 (NKp46) | BV650 | 740627 | BD Biosciences | Flow cytometry |
| Fas/CD95 | APC | BLE152604 | Ozyme | Flow cytometry |
| LLO pentamer GYKDGNEYI | PE | 178 | ProImmune | Flow cytometry |
| FLIP | - | 13250269 | ThermoFisher | Flow cytometry |
| Perforin | PE | BLE154306 | Ozyme | Flow cytometry |
| Granzyme B | Pacific blue | BLE372218 | Ozyme | Flow cytometry |
| CD127 | BV421 | BLE135023 | Ozyme | Flow cytometry |
| IFN$\gamma$ | BV711 | BLE505835 | Ozyme | Flow cytometry |
| KLRG1 | PE | BLE138408 | Ozyme | Flow cytometry |
| Cleaved caspase-3 biotinylated | - | 550557 | BD Biosciences | Flow cytometry |
| Streptavidin | APC | BLE405243 | Ozyme | Flow cytometry |
| CD11b | - | 550282 or 13-0112 | BD Biosciences | Microscopy |
| LY-6G | - | 551459 | BD Biosciences | Microscopy |
| LY-6C | - | Ab54223 | Abcam | Microscopy |
| Listeria | - | 294562 | Denka Seiken | Microscopy |
| c-Met | - | AF527 | R&D | Microscopy |
| Phospho-Akt | - | 05-1003 | Millipore | Microscopy |
| IgG Rat-Alexa Fluor A488 | AF488 | A11006 | ThermoFisher | Microscopy |
| IgG Rabbit-Alexa Fluor A546 | AF546 | A11035 | ThermoFisher | Microscopy |
| LAMP-1 | - | 553792 | BD Pharmigen | Microscopy |
| InlB | - | (*82* ) | | Western blot |
| Ef-tu | - | (*83* ) | | Western blot |
| IgG Rabbit-Peroxidase | - | A6154 | Sigma | Western blot |
